# Comparative transcriptomics of Hirschfeldia incana and relatives highlights differences in photosynthetic pathways

**DOI:** 10.1101/2023.10.18.562717

**Authors:** Francesco Garassino, Sofia Bengoa Luoni, Tommaso Cumerlato, Francisca Reyes Marquez, Jeremy Harbinson, Mark G. M. Aarts, Harm Nijveen, Sandra Smit

**Affiliations:** Laboratory of Genetics, Wageningen University & Research, Droevendaalsesteeg 1, 6708 PB, Wageningen, The Netherlands; Laboratory of Biophysics, Wageningen University & Research, Stippeneng 4, 6708 WE, Wageningen, The Netherlands; Bioinformatics Group, Wageningen University & Research, Droevendaalsesteeg 1, 6708 PB, Wageningen, The Netherlands

**Author notes:** Corresponding author: Sandra Smit.

## Abstract

Photosynthesis is the only yield-related trait that has not yet been substantially improved by plant breeding. The limited results of previous attempts to increase yield via improvement of photosynthetic pathways suggest that more knowledge is still needed to achieve this goal. To learn more about the genetic and physiological basis of high photosynthetic light-use efficiency (LUE) at high irradiance, we study *Hirschfeldia incana*. Here, we compare the transcriptomic response to high light of *H. incana* with that of three other members of the Brassicaceae, *Arabidopsis thaliana, Brassica rapa*, and *Brassica nigra*, which have a lower photosynthetic LUE.

First, we built a high-light, high-uniformity growing environment in a climate-controlled room. Plants grown in this system developed normally and showed no signs of stress during the whole growth period. Then we compared gene expression in low and high-light conditions across the four species, utilizing a panproteome to group homologous proteins efficiently. As expected, all species actively regulate genes related to the photosynthetic process. An in-depth analysis on the expression of genes involved in three key photosynthetic pathways revealed a general trend of lower gene expression in high-light conditions. However, *H. incana* distinguishes itself from the other species through higher expression of certain genes in these pathways, either through constitutive higher expression, as for *LHCB8*, ordinary differential expression, as for *PSBE*, or cumulative higher expression obtained by simultaneous expression of multiple gene copies, as seen for *LHCA6*.

These differentially expressed genes in photosynthetic pathways are interesting leads to further investigate the exact relationship between gene expression, protein abundance and turnover, and ultimately the LUE phenotype. In addition, we can also exclude thousands of genes from “explaining” the phenotype, because they do not show differential expression between both light conditions. Finally, we deliver a transcriptomic resource of plant species fully grown under, rather than briefly exposed to, a very high irradiance, supporting efforts to develop highly efficient photosynthesis in crop plants.

## Background

Considering the projected global population growth, the increasing effects of global warming, and the need for a more sustainable means of production, it is evident that the agricultural sector is under substantial pressure to increase crop yields while reducing land use and inputs such as fertilisers and pesticides. Over the past decade photosynthesis has taken a central role in plant research aimed at increasing crop yields because it plays a major role in the crop energy conversion efficiency, the only yield-related trait of food and feed crops that has not yet been maximised or even substantially improved by plant breeding [1].

While increasing crop productivity via improved photosynthetic efficiency was proposed over forty years ago [2], limited results have been achieved so far due to the physiological and genetic complexity of the photosynthetic process. Studies based on modelling of the photosynthetic process, bottleneck identification, and genetic modification aimed at overcoming identified bottlenecks have proven successful in some field crops, with yield increases ranging between 15 and 28% [3, 4, 5, 6, 7]. However, the inconsistency of results over multiple seasons [4] and across species [8] or growing conditions [9] indicates that more knowledge is needed on photosynthetic processes and how they are influenced by the environment across a range of timescales if we are to systematically increase the photosynthetic efficiency in crops.

Plant photosynthesis is defined as the process in which energy from light radiation is converted into chemical energy via a complex series of reactions resulting in the production of carbohydrates and oxygen [10]. A first set of photosynthetic reactions, catalyzed by photosystem complexes and an associated electron transport chain, is responsible for converting light radiation energy into chemical energy. This chemical energy is then stored in metabolically useful reducing agents (e.g. NADPH) and the energy-rich phosphate donor ATP. The processes linking light absorption to the formation of ATP and NADPH and other reducing agents collectively form the light reactions of photosynthesis [11]. The energy-rich re-ducing agents and ATP then drive a second set of photosynthetic reactions, the so-called dark reactions. These, generally referred as to the Calvin-Benson cycle, result in the conversion of the inorganic carbon substrate CO_2_ into organic carbohydrate molecules [11]. Bottlenecks or constraints - sites whose modification could result in improved photosynthesis - have been identified in both the light and dark reactions [1]. Furthermore, bottlenecks affecting photosynthesis have been identified in processes that would not be defined as strictly photosynthetic, such as the diffusion of CO_2_ into and through leaves to the site of CO_2_fixation in chloroplasts, and the transport of carbohydrates from photosynthetically active cells to carbon sinks elsewhere in the plant [12].

Our current knowledge of the key mechanisms and components of photosynthesis is the result of decades of studies in plants and other photosynthetic organisms [13]. This amounts to a vast body of knowledge, but on its own it is insufficient to improve crops’ photosynthesis and their yield. Studies conducted so far have highlighted how the link between crop photosynthesis and productivity is much more complex than originally thought, as a result of interactions of this process with plant development and environmental factors [14]. One characteristic of photosynthesis that can have a major impact on crop productivity is the decreasing light-use efficiency that occurs with increasing irradiance, giving rise to the light-saturation of photosynthesis and limitation of assimilation rate. This limitation has a substantial impact on productivity at irradiance levels normally recorded during summer in temperate areas of our planet. We define photosynthetic light-use efficiency (LUE) as the ratio between photosynthetically assimilated CO_2_ and incident light radiation, or irradiance. The decrease in LUE due to increasing irradiance is well-known and its causes are linked to both limitations in the photosynthetic process and other associated processes [15].

Evidence has been reported for large natural variation in photosynthesis rates, and therefore photosynthetic LUE, among crop and other plant species [16, 17]. This suggests that a degree of plasticity exists for photosynthesis that could be leveraged to increase the photosynthesis of crop species. However, it is nowadays clear that increases might only be achieved if knowledge is accumulated on the regulation of the photosynthetic process as well as specific strategies some plant species might have evolved that result in photosynthesis optimised to meet unusual goals [15]. One powerful way to map the genetic basis of complex biological processes is via the analysis of the associated transcriptional activity. Over the past years, several studies on transcriptional activity in a number of species have increased our knowledge on the response of photosynthesis to irradiances of different intensities or changes in irradiance. It was shown that *Arabidopsis thaliana* acclimates to high light by increasing expression of heat shock response genes and lipid remodelling genes [18], that rice (*Oryza sativa*) exposed to variations in irradiance associated to field conditions activates a large number of biotic and abiotic stress genes [19], and that barley (*Hordeum vulgare*) expresses genes involved in phenolic compounds accumulation at a higher level with increasing irradiance [20]. However, none of these studies applied a long-term, very high irradiance treatment, similar to what is experienced by plants growing in natural temperate environments during summer months, at high altitude conditions, or in the equatorial region. Neither did they include species with a particularly high photosynthetic LUE. We consider these two factors essential for unraveling the physiological and genetic basis of photosynthetic light-use efficiency, and for ultimately building more light-use efficient photosynthesis in our crops [15].

Here, we present the analysis of gene expression in *Hirschfeldia incana* (L.) Lagr.-Foss., the species we previously proposed as preferred model for studies on high photosynthetic LUE [21, 15]. The gene expression under contrasting high- and low-light irradiance conditions is compared to that of Brassicaceae family relatives *A. thaliana, Brassica rapa*, and *Brassica nigra*. While *A. thaliana* does not share the whole genome triplication that the other three species underwent and is therefore more distantly related, *B. rapa* and *B. nigra* represent the different evolutionary history of two major lineages emerging after this event [22]. Using transcriptomics we aim to elucidate which genes and thus pathways are involved in the maintenance of a high photosynthetic LUE at high irradiance in *H. incana*. First, we describe the experiment we conducted under high irradiance, and present the results of differential gene expression (DGE) analysis performed on each of the four species. Then, we report on the use of a panproteome to compare gene expression changes across the four species, and on the exploration of common and divergent trends in the gene expression response to high-light by means of untargeted enrichment analyses. We then present the results of targeted analysis of expression patterns across the four species for genes involved in key photosynthesis-related pathways. Lastly, we discuss the findings in the light of their implications for *H. incana’s* higher photosynthetic light-use efficiency at high irradiance. Our work thus describes the transcriptional differences associated with plant growth under highly contrasting irradiance conditions, and serves as a resource for the elucidation of the genetic determinants of the striking photosynthetic capacity of *Hirschfeldia incana*.

## Materials and Methods

### Construction of high-uniformity growth system

Two custom light ceilings were built for this study. Each ceiling measured approximately 4.3 m^2^ (*l* 175 cm, *w* 245 cm), was equipped with six dimmable VYPR2p LED fixtures (Fluence, Austin, USA) arranged in three equally spaced rows (between-rows distance of 60 cm, and was hung so that fixtures would be one meter high over plants. We then centered two custom-made growing tables measuring approximately 1.6 m^2^ (*l* 118 cm, *w* 137 cm) under the custom light ceilings, divided each table into thirty growing areas, each measuring approximately 0.05 m^2^, and performed irradiance measurements at the centre of each growing area. By calculating averages over the thirty areas under each light ceiling, we optimised the output of the LED fixtures to have average irradiances as close as the reference values we chose for our treatments.

### Plant material and growing conditions

*Hirschfeldia incana* accession ‘Nijmegen’, *Brassica nigra* accession ‘DG1’, *Brassica rapa* R-o-18, and *Arabidopsis thaliana* Col-0 were used for this experiment. ‘Nijmegen’ is an inbred line (over six rounds of inbreeding) from an *H. incana* plant originally collected in Nijmegen, The Netherlands; ‘DG1’ is a second-generation inbreeding line of *B. nigra* sampled from a natural population near Wageningen, The Netherlands; and ‘R-o-18’ is a *B. rapa* inbred line [23, 24].

Seeds of all species were germinated on a peat-based potting mix for nine days under an irradiance of 200 µmol m^*-*2^ s^*-*1^. Twelve healthy seedlings per species were then transferred to 2 L pots (ϕ 13.9 cm, h 17.4 cm, Soparco, Condé-sur-Huisne, France) filled with a peat-based potting mixture enriched with perlite and 2.5 g*/*L Osmocote® Exact Standard 5-6M slow-release fertiliser (ICL Specialty Fertilizers, Geldermalsen, The Netherlands).

Plants were germinated and grown in a climate-controlled room equipped with the custom arrays of high-output LED light modules described above, with a photoperiod of 12 h day and 12 h night, and air temperature set at 23 ^*0*^C and 20 ^*0*^C, respectively. Humidity and CO_2_ levels were set at 70% and 400 ppm. Six plants per species were assigned to the high light (HL) treatment of 1800 µmol m^*-*2^ s^*-*1^ (measured irradiance average: 1843.6 µmol m^*-*2^ s^*-*1^) and the remaining six to the low light (LL) treatment of 200 µmol m^*-*2^ s^*-*1^ (measured irradiance average: 227.5 µmol m^*-*2^ s^*-*1^). Ir-radiance uniformity was very high for both HL and LL treatments, with a U_2_ value (defined as *minimum ir-radiance/maximum irradiance*, [25, 26]) of 0.93. Plant positions were randomised across growing areas. Plants assigned to the LL treatment were fertigated daily, while plants assigned to the HL treatment were fertigated twice a day, with a custom nutrient solution (0.6 mM NH_4_^+^, 3.6 mM K^+^, 2 mM Ca^2+^, 0.91 mM Mg^2+^, 6.2 mM NO_3_^−^,1.66 mM SO_4_^2–^, 0.5 mM P, 35 µM Fe^3+^, 8 µM Mn^2+^, 5 µM Zn^2+^, 20 µM B, 0.5 µM Cu^2+^, 0.5 µM Mo^4+^).

### Sampling and RNA extraction

Twenty-eight days after sowing, samples representative of the whole canopy were collected from all plants. All leaves (for smaller plants such as *A. thaliana* and *H. incana*, especially when grown under low light) or half the total number of leaves were excised from plants, transferred to 50 mL tubes, and flash-frozen in liquid nitrogen. All leaf samples were subsequently crushed with a mortar and pestle in excess liquid nitrogen, and further homogenised with glass beads for 2 min at 30 Hz in a MM300 Mixer Mill (Retsch GmbH, Haan, Germany). Total RNA was extracted with the RNeasy Plant Mini Kit (QIAGEN N.V., Venlo, The Netherlands) according to manufacturer’s instructions, and eluted using 50 µL of DNAse/RNAse-free water. The following DNAse treatment and RNA recovery were performed as described in [27]. 6 µL of 10X DNAse buffer and 4 µL of RQ1 DNAse (Promega, Leiden, The Netherlands) were added to 50 µL of RNA, and incubated for 30 minutes at 37 ^*0*^C.

The RNA was then precipitated overnight using ammonium acetate and ethanol, and resuspended in 25 µL of DNAse/RNAse-free water. To check RNA quality and integrity, 1 µL of RNA was used to (1) load a 1% agarose-Ethidium bromide gel and after electrophoresis observe the bands using standard imaging and (2) to determine spectrophotometric parameters with a Nanodrop 2000 (Thermo Fisher Scientific Inc., Waltham, U.S.A.). The RNA was further quantified using the Qubit RNA BR Assay kit and a Qubit 4 fluorometer (Thermo Fisher Scientific Inc., Waltham, U.S.A.).

### Sequencing

RNA from five of the six plants of each species grown under each light treatment was sequenced by Novogene (UK) Company Ltd., Cambridge, U.K.. Poly-A enriched RNA was employed to prepare sequencing libraries with the NEBNext® Ultra™ RNA Library Prep Kit (New England Biolabs, Ipswich, U.S.A.). Paired-end, 150-bp-long reads (PE150) were generated with a NovaSeq 6000 system (Illumina Inc., San Diego, U.S.A.) aiming at obtaining 6 Gb of data per sample.

### Selection and preparation of genome assemblies and annotations

For mapping of sequencing reads and quantification of gene expression, the TAIR10 [28] genome assembly and the Araport11 annotation [29] were used for *A. thaliana*, the “Chiifu” v3.0 assembly and annotation [30] were used for *B. rapa*, the “Ni100” v2.0 assembly and annotation [31] were used for *B. nigra*, and the “NIJ6” v1.0 assembly and annotation [21] were used for *H. incana*. For pan-proteome building, the v3.0/3.1 *A. arabicum* [32], the v1.0 *R. raphanistrum* [33], the v1.0 *R. sativus* [34], and the v1.0 *S. irio* [35] genome assemblies and annotations were employed together with the aforementioned ones. The exact locations where the various files were downloaded from can be found in Table S13.

Statistics were collected for all genome assemblies and annotations with custom Python (v3.11.0) scripts and are reported in Table S14. Given that not all genome annotations contained multiple transcript isoforms, all GFF files were processed with the agat_sp_keep_longest isoform.pl script from the AGAT toolkit v1.0.0 [36] to generate annotations containing only the longest transcript isoforms of all gene models. Subsequently, these GFF files were filtered with the agat_sp_filter_by_ORF_size.pl script to remove all gene models that would have yielded protein sequences shorter than 30 amino acids. Finally, a number of gene models identified in the *R. raphanistrum* and *R. sativus* annotation that would still not result in protein sequences (due to stop codons embedded in their sequence) were removed from the corresponding annotations with the agat_sp_filter_feature_from_kill_list.pl script. The resulting filtered annotation files are provided with the data package linked to this article.

### Identification and Analysis of Differentially Expressed Genes

The quality of sequencing libraries was assessed with MultiQC [37] v1.11. A snakemake (v7.19.1) [38] pipeline was employed to automate subsequent read mapping and transcript quantification steps. Reads were aligned to reference genome assemblies with two passes of the STAR [39] v2.7.10a aligner (STAR indexing running with parameters -sjdbOverhang 139 and – genomeSAindexNbases 13, STAR aligner running with parameter –clip5pNbases 10 10). Assembly and quantification of full-length transcripts were then achieved with StringTie [40] v2.2.1 (running with option -e). Per-sample gene and transcripts counts were then grouped by species with the prepDE Python script included in the StringTie suite (running with parameter -l 140). Transcripts per million (TPM) counts [41] were extracted for visualisation purposes from the StringTie output with a custom Python script.

Relationships between samples of the same species were explored with PCA plots of transcript counts transformed by means of regularized logarithm [42]. Differen-tially Expressed Genes (DEGs) were subsequently identified with DESeq2 [42] v1.34.0 running in R [43] v4.1.1.

### Panproteome construction

Proteomes were created from the filtered annotations of all eight species with the AGAT toolkit agat sp extract sequences.pl script, running with options -p, –cis, and –cfs. A panproteome was subsequently constructed by running PanTools v4.1.0 [44, 45] commands build panproteome, busco protein, (with options-if brassicales odb10 –version busco4), optimal grouping, and change grouping (with option –version 4, and thus running with a relaxation parameter of 4). A separate panproteome was constructed featuring chloroplast proteomes for *A. thaliana, B. rapa, B. nigra*, and *H. incana* with PanTools commands build panproteome and group (with the same relaxation parameter of 4). The panproteome was visualized by making UpSet plots [46] with the ComplexUpset package (v1.3.3) running in R v4.4.2.

### Integration of panproteome and DE results

The homology table resulting from panproteome construction was integrated with Differential Expression analysis results by means of a custom script running in Python v3.10.9 and leveraging NumPY v1.24.1 [47], and Pandas v1.5.3 [48]. The resulting homology/DE status table was further processed and visualised with a custom script running in R v4.2.2. A heatmap of non-ambiguously responding core groups was generated with the Pheatmap v1.0.12 package. After specific categories of homology groups were selected, a Gene Ontology (GO) Biological Process (BP) enrichment analysis was performed for the *A. thaliana* gene identifiers present in said groups with TopGO v2.50.0 [49], relying on the org.At.tair.db v3.16.0 Bioconductor annotation data package, running the “Classic” algorithm, and performing Fisher tests. Enrichment results for each set of groups were filtered by keeping only terms which were associated to at least five genes. A KEGG pathway enrichment analysis was subsequently performed on the *A. thaliana* genes present in the same categories of homology groups with the enrichKEGG function of the Clus-terProfiler v4.6.2 package [50, 51]. For both enrichment analyses, the set of background genes (i.e., the analysis “universe”) was composed by all *A. thaliana* genes surviving the DE analysis (i.e., genes for which an adjusted *p*-value could be calculated by DESeq2).

### Processing and visualization targeted analysis results

Expression profiles, TPM-normalized counts and homology relationships were processed and visualized with custom R scripts making use of packages dplyr (v1.1.0), ggplot2 (v3.4.1), janitor (v2.2.0), pheatmap (v1.0.12), scales(v1.2.1), stringr (v1.5.0), tidyr (v1.3.0). All scripts are available at https://doi.org/10.4121/5b88cdf2-eb5f-4033-8ece-1f3f488a1f83.

## Results

### Plant growth under a reliable high-light environment

In this study, we aimed to identify genes and pathways responsible for the higher photosynthetic light-use efficiency of *H. incana* under high-irradiance conditions. To create strongly contrasting growth conditions, we set our low light (LL) irradiance to 200 µmol m^*-*2^ s^*-*1^ and our high light (HL) irradiance to 1800 µmol m^*-*2^ s^*-*1^ for 12 h per day. We calculated U_2_ irradiance uniformity values [25, 26] over all growing positions designated for both treatments, and we selected positions on each growing table that resulted in the best irradiance uniformity. For the LL table, this resulted in an average irradiance of 227.5 µmol m^*-*2^ s^*-*1^ associated with an U_2_ of 0.93, while for the whole HL table we measured an average irradiance of 1843.6 µmol m^*-*2^ s^*-*1^, also associated with an U_2_ of 0.93.

To compare the light treatments to conditions that plants would experience in natural environments, we calculated a Daily Light Integral (DLI) [52], a measure of the total irradiance delivered over the course of a day per unit of area, for each treatment. This resulted in DLIs of 9.82 mol m^*-*2^ d^*-*1^ and 79.64 mol m^*-*2^ d^*-*1^ for the LL and HL treatments, respectively.

Besides *H. incana*, this study featured three other Brassicaceae species: *A. thaliana, B. rapa*, and *B. nigra*. These are the same we used for previous work in which they showed to have lower photosynthetic LUE than *H. incana* [21]. Plants of the four species established and grew well under both light treatments, albeit with differences in growth and architecture (Figures 1, S1). No stress symptoms were visible on the plants throughout the growing period. 20% of the *B. nigra* plants from the LL treatment appeared to grow more slowly and had paler leaf color than the other *B. nigra* plants (Figure S1).

**Figure 1:**
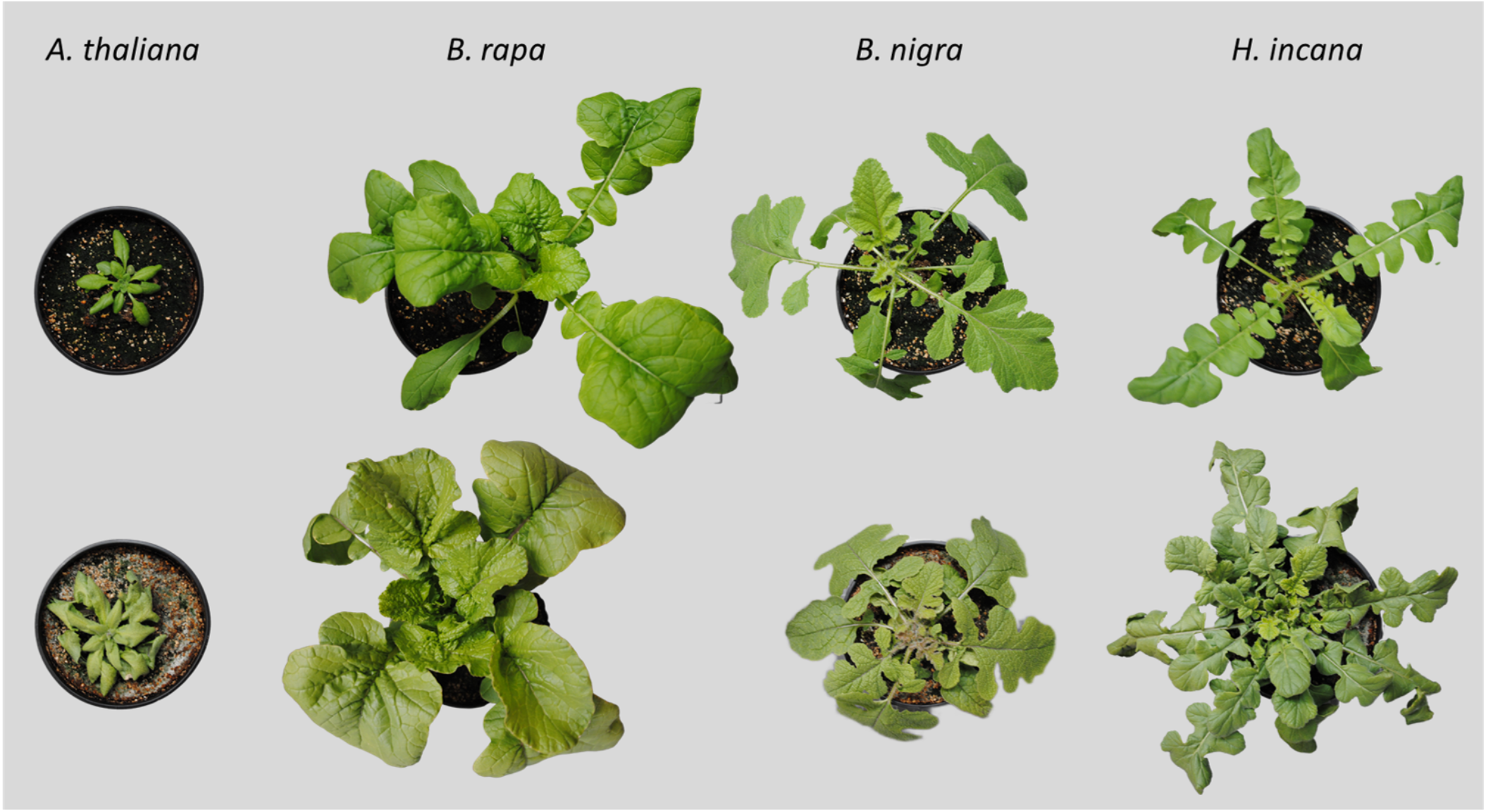
Pictures of representative plants at the end of the experiment for each of the four species grown under the two irradiance conditions. Left to right: *A. thaliana, B. rapa, B. nigra, H. incana*. Top row: low light (LL) irradiance. Bottom row: high light (HL) irradiance.

### Per-species differential gene expression analysis

To study the gene expression under contrasting light conditions in the four species we sequenced forty mRNA libraries (4 species ×5 replicates × 2 conditions) generated from RNA extracted from whole plant canopies, with an average of 22.2 ± 2.4 million reads per library. The MultiQC inspection of all sequencing reads did not show any quality issues in our dataset. Percentages of reads mapped to reference genomes were high, ranging between 93.4 ± 1.5% and 96.3 ± 0.7% (Table S1).

We performed differential expression (DE) analysis on data from each species individually with DESeq2 and selected all differentially expressed genes (Tables 1, S2-S5). Per-species principal component analysis performed on regularized logarithm-transformed count data showed that the general patterns of gene expression are consistent across biological replicates belonging to the same species and originating from the same treatment (Figure S2). The percentages of genes significantly differentially expressed were similar for *A. thaliana, B. rapa*, and *H. incana*, while they were lower for *B. nigra* due to the high number of genes in the annotation. Since we are interested in differences and similarities between *A. thaliana* and the other species, and in particular *H. incana*, we compared the gene expression across the species.

**Table 1.** Numbers of differentially expressed and non-deferentially expressed genes for the various species. Percentages of the total number of genes are placed between brackets. Significant differences for *p<*0.05.

|  | Number of genes |  |  |
| --- | --- | --- | --- |
|  | Higher expression | Lower expression | Unchanged expression |
| *A. thaliana* | 3,346<br>(12.1%) | 3,027<br>(11.0%) | 21,226<br>(76.9%) |
| *B. rapa* | 7,292<br>(15.7%) | 7,138<br>(15.4%) | 32,050<br>(68.9%) |
| *B. nigra* | 4,723<br>(7.9%) | 4,052<br>(6.8%) | 50,934<br>(85.3%) |
| *H. incana* | 4,284<br>(13.2%) | 4,334<br>(13.3%) | 23,900<br>(73.5%) |

### Cross-species comparison using a panproteome

To enable the comparison of gene expression across species, we built a panproteome to group homologous genes (orthologs and paralogs) [44]. A panproteome of eight Brassicaceae species (*Aethionema arabicum, A. thaliana, B. nigra, B. rapa, H. incana, Raphanus raphanistrum, Raphanus sativus, Sisymbrium irio*) yielded 106,511 homology groups (HGs, Figure S3, Table S6). We then selected HGs containing at least one gene from one of the four species for which RNA-Seq was performed, leaving 63,675 HGs for downstream analysis (Figure 2).

**Figure 2:**
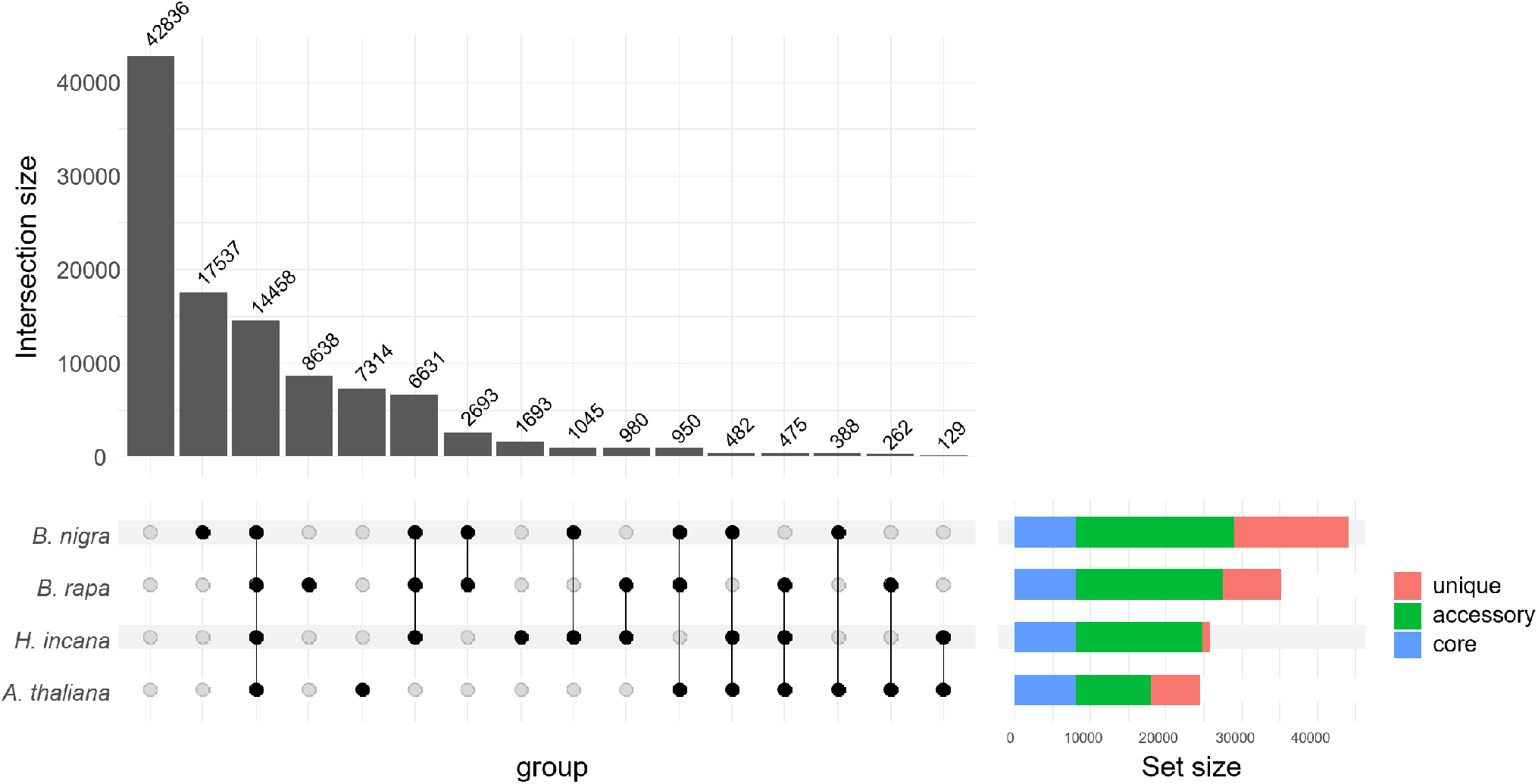
An UpSet plot of the panproteome homology groups (HGs), based on the four species for which RNA-Seq was done. Vertical bars represent the number of HGs, classified by presence/absence of genes from the various species as illustrated at the bottom of the figure. The first bar represents all the HGs of the pan-proteome constructed with proteomes from eight Brassicaceae species that do not contain any genes from the four species we sequenced. The horizontal colored bars show how many of the HGs containing at least one gene from a species fall within one of three categories: core HGs, i.e. those containing at least one gene from all four species; accessory HGs, i.e. those containing genes from more than one, but not all, species; and unique HGs, i.e. those containing only genes from a single species.

We also distinguished “differentially expressed” (DE) HGs, which contain at least one gene that was differentially expressed between both light conditions, and “non-DE” HGs (Table 2), which do not. Among the 19,012 DE HGs, the 10,770 which have an ortholog from each of the four species (i.e. the core proteome) form the main target of our research. Of particular interest are the differences and similarities between *A. thaliana* and the other species, all of which are members of tribe Brassiceae, and in particular *H. incana*. Similarly, we identified 3,688 HGs which contain genes that were not differentially expressed in any of the species and can therefore not explain the phenotypic differences (Table S7). Based on the *A. thaliana* genes contained in these groups, we found 267 Gene Ontology (GO) Biological Process (BP) terms enriched (Table S8). As expected, no terms related to photosynthesis or high-light adaptation were identified in this set of groups.

**Table 2.**
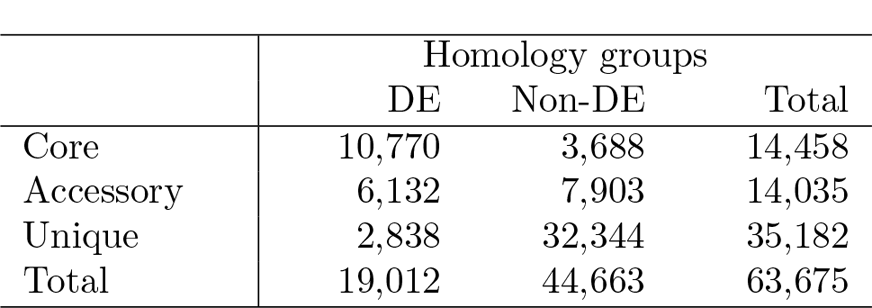
Numbers of homology groups (HGs) in the constructed panproteome, classified based on their “differential expression” (DE) status. An HG is classified as “DE” if it contains at least one gene differentially expressed between both light conditions. If this condition is not met, the HG is classified as “non-DE”.

|  | Homology groups |  |  |
| --- | --- | --- | --- |
|  | DE | Non-DE | Total |
| Core | 10,770 | 3,688 | 14,458 |
| Accessory | 6,132 | 7,903 | 14,035 |
| Unique | 2,838 | 32,344 | 35,182 |
| Total | 19,012 | 44,663 | 63,675 |

To compare transcriptional activity across species within a single light condition, we compared transcriptsper-million (TPM)-normalised transcript counts. To assess bias due to differences in sequencing libraries, which are not corrected for during TPM normalisation [53], we tested whether average transcript abundances were similar across species. We selected the non-DE homology groups containing a single expressed ortholog for each of the four species. We averaged TPM counts (regardless of treatment) for each species and calculated per-HG log_2_-ratios between average counts for the single orthologs of the various species. The distribution of these ratios showed that on average the *A. thaliana* transcript abundances are higher than those of *B. rapa* and *H. incana* (% of area under the curve (AUC) for log_2_-ratios *>* 1: 59.9 and 62.8%, respectively), which are in turn higher than those of *B. nigra* (% of area under the curve for log_2_-ratios *>* 1: 67.4 % for *B. rapa* and 67.2% for *H. incana*) (Figure S4, Table S9). Given the fact that expression in *H. incana* is generally lower than in *A. thaliana* and similar to that in *B. rapa*, we conclude that detection of a significantly higher expression in *H. incana* is the effect of biological processes rather than an artifact.

### Comparative analysis of core DE homology groups highlights photosynthetic pathways

Of the 10,770 core HGs (CHGs) containing at least one gene differentially expressed under HL, 10,352 showed non-ambiguous differential expression within each species and were selected for downstream analysis. We defined non-ambiguous DE as the situation in which the expression of all genes is exclusively increased or decreased.

Clustering the CHGs with non-ambiguous responses allowed us to identify expression profiles for the four species (Figure 3, Table S10). Some CHGs show consistent higher or lower expression in all species (245 and 382, respectively). More often, an higher or lower expression is shared by some species (2,165 and 1,890 CHGs) or is unique to a species (2,163 and 1,946). Lastly, there are CHGs showing contrasting expression (higher expression in some species, lower expression in others) across species (1,561).

**Figure 3:**
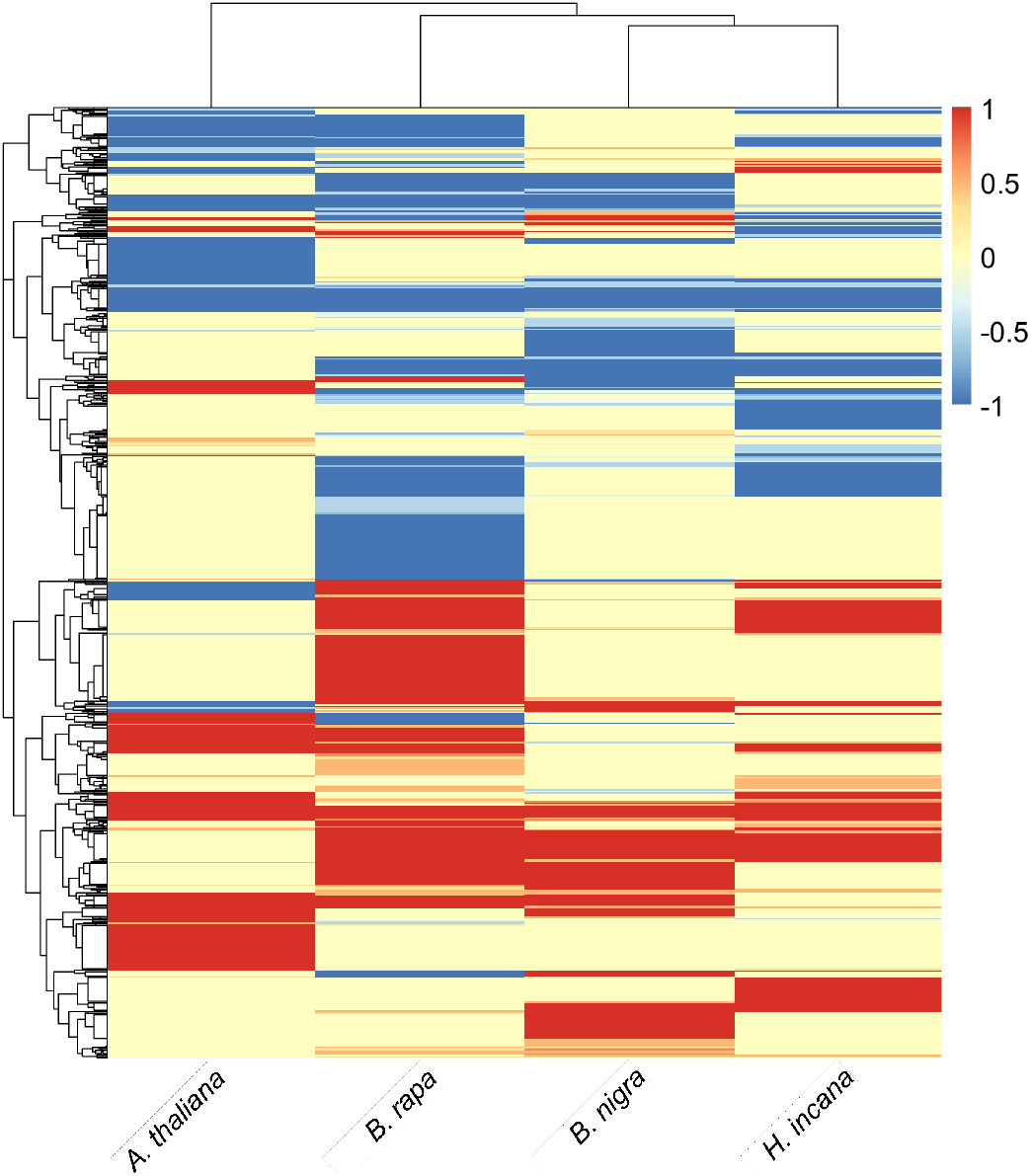
Heatmap of 10,352 groups showing only non-ambiguous responses per species. The color scale represents the ratio between the number of higher (positive numbers) or significantly lower expressed (negative numbers) genes and the total number of gene models present in each group per each species. Both rows and columns were clustered with hierarchical clustering based on Euclidean distances.

In order to get an overview of the role of the genes belonging to clusters of CHGs, we performed Gene Ontology (GO) [54, 55] and KEGG [56, 57] enrichment analyses separately for clusters of CHGs containing at least one gene model with significantly higher or lower expression in each species, in the three Brassiceae species (*B. rapa, B. nigra, H. incana*), and in *H. incana* alone. GO enrichment analysis for the sets of *A. thaliana* genes in the clusters of CHGs containing genes with higher expression under HL resulted in terms related to response to water deprivation and salt stress, heat, low cellular oxygen, and flavonoid biosynthesis (Table S11). A similar selection and analysis for genes with lower expression under HL resulted in terms involved in high-light response, chlorophyll metabolism, and growth regulation.

KEGG enrichment analysis highlighted seventeen over-represented pathways (Table S12). Twelve pathways were enriched in CHGs containing at least one gene with higher expression under HL in all species, Brassiceae species, or *H. incana* alone. The most notable of these twelve pathways was “Carbon metabolism” (ath01200). The remaining five patwhays were enriched in CHGs containing at least one gene with lower expression under HL in all species, Brassiceae species, or *H. incana* alone. Notably, these five pathways comprised the two currently annotated in KEGG for photosynthesis: “Photosynthesis”(ath00195), and “Photo-synthesis - antenna proteins” (ath00196). Since all three photosynthesis-related KEGG pathways were highlighted by our enrichment analysis, we decided to further explore the expression of the genes associated with these pathways in search of clues on the higher photosynthetic LUE of *H. incana*.

### Targeted analysis of light-harvesting complex genes

We first analyzed the expression patterns of the *A. thaliana* genes annotated with the KEGG pathway “Pho-tosynthesis - antenna proteins” (ath00196) and their orthologs in *B. rapa, B. nigra*, and *H. incana*. This allowed us to investigate transcriptional differences associated with light-harvesting complexes (LHCs), which are amongst the first complexes involved in the photosynthetic process. The KEGG pathway is made up of 22 *A. thaliana* genes assigned to 14 homology groups. These groups contain 34 genes for *B. rapa*, 33 genes for *B. nigra*, and 35 genes for *H. incana*. Inspection on these genes revealed log_2_ fold change (log_2_FC) values ranging between -3.15 and 1.01, with almost all genes showing significant lower expression under HL, except for *LHCB8, LHCB7*, and *LHCA5* (Figure S5). No differences across species were observed except for two genes coding for photosystem II (PSII) antenna proteins, *LHCB8* and *LHCB7*, and two coding for photosystem I (PSI) antenna proteins, *LHCA6* and *LHCA5* (Figure 4a). Considering the particular features explained below, we selected the *LHCB8* and *LHCA6* genes for further investigation.

**Figure 4:**
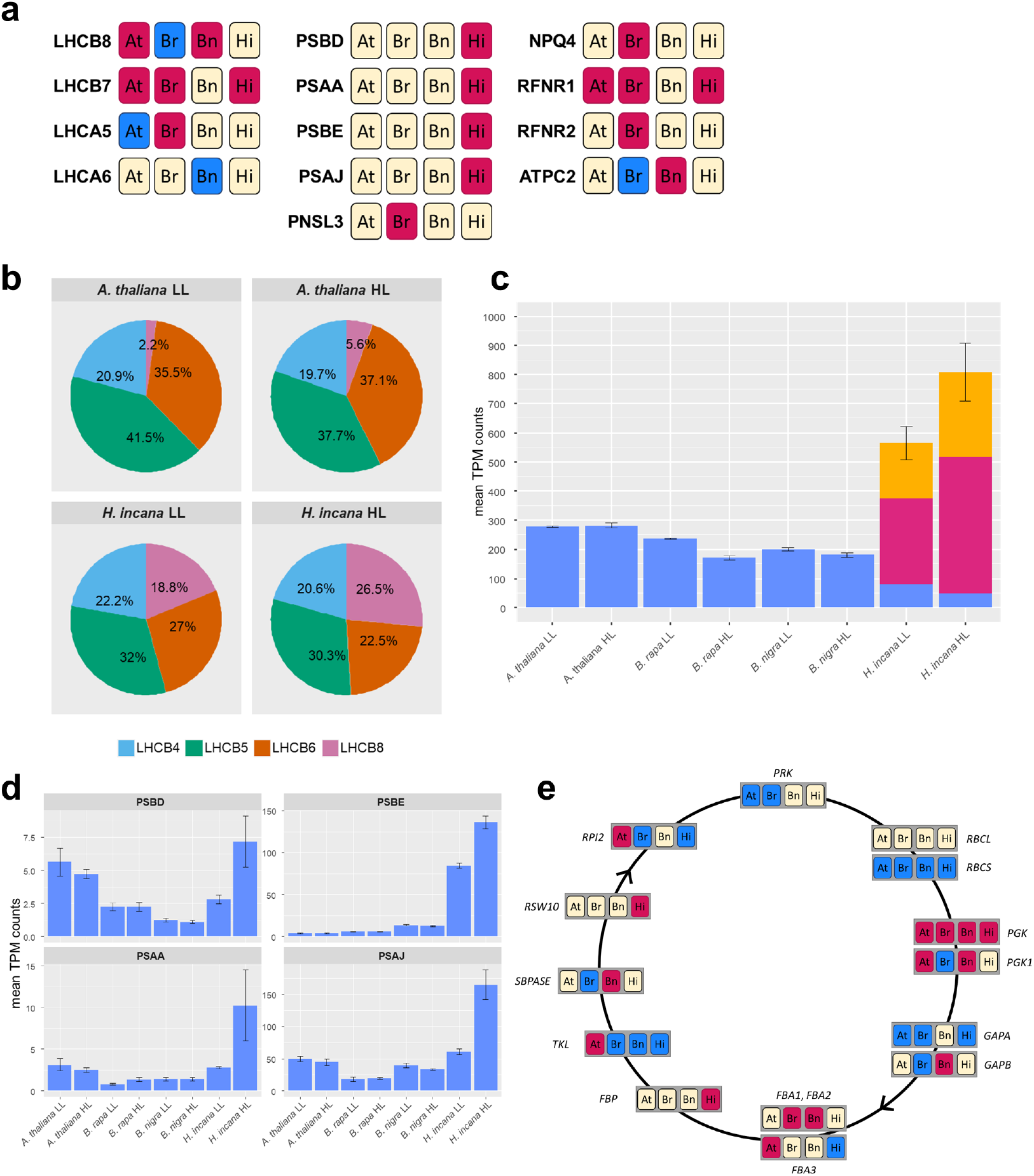
Results of the targeted analysis of photosynthetic pathways. (a) Schematic view of the restricted set of genes belonging to KEGG pathways ath00196 (“Photosynthesis - antenna proteins”, left column) and ath00195 (“Photosynthesis”, center and right columns) showing higher expression under HL in at least one of the species. Magenta box indicate significantly higher expression under HL, while blue boxes indicate significantly lower expression under HL. At: *A. thaliana*; Br: *B. rapa*; Bn: *B. nigra*; Hi: *H. incana*. (b) Pie charts representing the relative abundance of transcripts originating from genes encoding components of the PSII minor antenna. (c) Mean normalized abundance of *LHCA6* transcripts in plants of the four species grown under the two irradiance treatments. The different colors represent different paralogs. Error bars represent the standard errors of the mean. The full comparison of transcript abundances for *LHCA6* can be found in Figure S7. (d) Mean normalized abundances of *PSBE, PSBD, PSAA*, and *PSAJ* transcripts in plants of the four species grown under the two irradiance treatments. Error bars represent the standard errors of the mean. Full across-species comparisons can be found in Figures S9, S10, S11, and S12. (e) Schematic representation of the Calvin-Benson cycle, or the “dark reactions” of photosynthesis, and differential expression status of genes involved in the four species. Yellow: no significant differential expression under HL; red: significantly higher expression under HL; blue: significantly lower expression under HL. At: *A. thaliana*; Br: *B. rapa*; Bn: *B. nigra*; Hi: *H. incana*.

*LHCB8* was first investigated as member of a subset of rarely expressed light-harvesting complex (LHC) protein encoding genes [58]. The LHCB8 protein resembles the CP29.1 and CP29.2 proteins, encoded in *A. thaliana* by the *LHCB4*.*1* and *LHCB4*.*2* genes, and is therefore also known as CP29.3 (*LHCB4*.*3*). *AtLHCB8* shows a different expression pattern than *AtLHCB4*.*1* and *AtLHCB4*.*2*, suggesting a different role for the protein in the LHC. It seems to be present only in species of the eurosids, a subclade of the rosids [58]. The LHCB8 protein is present as a monomer within the PSII supercomplex, forming the so-called “minor antenna” of photosystem II with a number of other LHCB proteins (LHCB4, LHCB5, LHCB6) [59]). The expression of *A. thaliana LHCB8* is induced by high-irradiance conditions [60]). *LHCB8* is a single-copy gene in the four species used for this study and under HL had moderately higher expression in *A. thaliana* and *B. nigra*, while it had moderarely lower expression in *B. rapa* and showed no significant changes in *H. incana*. Based on transcripts-per-million (TPM)-normalised read counts, *LHCB8* transcripts represent roughly 26.5% of the transcript pool for the minor antenna in *H. incana* plants grown under HL, while they represent only 5.6%, in *A. thaliana* (Figure 4b). This high representation is also found in *B. rapa* and *B. nigra*, with *LHCB8* making up 22.5% and 31.1% of minor antenna transcripts (Figure S6).

*LHCA6* is a poorly-expressed gene coding for a protein associated with PSI as an antenna monomer. *LHCA6* is present as a single-copy gene in *A. thaliana, B. rapa*, and *B. nigra*, while it has three tandem copies in *H. incana* [21]). *LHCA6* did not show statistically significant changes in expression in *B. nigra, A. thaliana* and in any of the three copies of *H. incana*. However, it had lower expression in *B. rapa* under HL. To determine expression across species, we summed TPM-normalized counts for the three *LHCA6* copies in *H. incana* and calculated all pairwise ratios between counts in the four species under the two treatments. Inspection of ratios between counts in *H. incana* and other species revealed the *LHCA6* paralogs to have higher expression in *H. incana* under both the irradiance conditions after correction for the transcriptional baseline differences (Figure 4c, S7).

### Targeted analysis of light reactions genes

We then analyzed the expression patterns for the CHGs containing the 77 *A. thaliana* nuclear and chloroplast genes that are annotated with the KEGG pathway “Photosynthesis” (ath00195), which are involved in the light reactions of photosynthesis. Log_2_FC values ranged between 1.98 and -1.78, with 142 of the total 368 genes significantly DE under HL. When considering only the significant expression changes, the trend across “Photosynthesis” pathway genes is lower expression under HL in all species: 29 genes out of 33 in *A. thaliana*, 48 out of 52 in *B. rapa*, 25 out of 26 in *B. nigra*, and 27 out of 31 in *H. incana* (FigureS8). A small number of CHGs contained at least one gene having higher expression under HL in one of the species (Figure 4a). For further analysis, we focused on those showing higher expression only in *H. incana*, thus selecting genes *PSBD* and *PSBE*, part of the PSII complex, and *PSAA* and *PSAJ*, part of the PSI complex.

The D2 protein, encoded by *PSBD*, forms the core of PSII along with the D1 protein, encoded by *PSBA*. These two subunits together bind three macromolecules that are fundamental for photosynthetic light reactions: the P680 reaction center, which transfers energy to water molecules, the Mn_4_CaO_5_ cluster responsible for the splitting of water molecules and retrieval of electrons, and components of the primary electron transfer chain, such as plastoquinones Q_A_ and Q_B_ [61]. The PSII reaction center is completed by the subunit encoded by the *PSBI* gene and cytochrome b559, composed of subunits encoded by the *PSBE* and *PSBF* genes and a heme co-factor [62]. The *PSBD* gene is highly expressed in *A. thaliana* plants grown under both treatments, and *H. incana* plants grown under HL, while the *PSBE* gene is highly expressed only in *H. incana* plants grown under HL (Figures 4d, S9, S10).

The photosystem I (PSI) core is composed of proteins encoded by the *PSAA* and *PSAB* genes. The PSI complex is composed of several additional subunits, including one stabilized by the protein encoded by gene *PSAJ* [63]. The expression of *PSAA* and *PSAJ* orthologs appears to be significantly higher in *H. incana* plants grown under HL, with plants growing under LL having similar transcript levels to those measured in the other species irrespective of the treatment (Figures 4d, S11, S12).

### Targeted analysis of carbon metabolism genes

Continuing our analysis based on photosynthesis KEGG-related pathways, we studied the expression of the 273 *A. thaliana* nuclear and chloroplast genes associated to the KEGG pathway “Carbon metabolism” (ath01200) and their orthologs (Figure S13). Pathway ath01200 comprises genes involved in both catabolism and anabolism of carbon-based molecules, organized in a number of modules. Inspection of these modules revealed that the genes related to the Calvin-Benson cycle, and thus to assimilation of inorganic carbon into the end product of photosynthetic reactions, carbohydrates, were grouped into module “Reductive pentose phosphate cycle (Calvin cycle)” (ath M00165). The expression of genes included in this module did not show an obvious profile (Figure S14). However, two of the 23 CHGs associated with this module contained genes that had higher expression uniquely in *H. incana* under HL. These are orthologs of the *A. thaliana* genes *FBP* and *RSW10* (Figure 4e). *A. thaliana* mutants for the *RSW10* gene has been linked to ribose-5-phosphate metabolism and cellulose biosynthesis, but no direct involvement with photosynthetic activity has been described to date [64, 65]. Gene *FBP*, instead, has been associated with photosynthetic activity, and *FBP* overexpression has been proven to increase soluble sugar and starch contents, as well as photosynthetic CO_2_ assimilation [66].

## Discussion

In this study, we explored the transcriptomes of plants of four Brassicaceae species (*A. thaliana, B. rapa, B. nigra, H. incana*) grown under contrasting irradiances to unravel the genetic determinants of *H. incana*’s high photosynthetic light-use efficiency under high irradiance. Considering the complexity of our dataset and based on the results of our untargeted enrichment analysis, we decided to restrict our exploration by focusing on genes related to photosynthesis in the KEGG ontology.

### Increasing power of elimination through super-natural irradiance

To make sure we would observe any transcriptional differences associated with growth under high-irradiance conditions, we designed and built a high-output, high-uniformity lighting system. The per-treatment daily light integral (DLI) of 9.82 mol m^*-*2^ d^*-*1^ that we measured for the low light (LL - 227.5 µmol m^*-*2^ s^*-*1^) treatment is consistent with what has been reported for winter months in warm-temperate climate areas, while the DLI of 79.64 mol m^*-*2^ d^*-*1^ we measured for the high light treament (HL - 1843.6 µmol m^*-*2^ s^*-*1^) is substantially higher than the values of 60-65 mol m^*-*2^ d^*-*1^ reported for summer months in the same climate areas [67, 52, 68, 69].

Our study differs from previous studies on high light responses not only because of our use of the “super-natural” magnitude of our high light treatment, but also for the way the treatment was applied. While previous high-light studies involving *A. thaliana* have employed irradiances ranging from between 150 and 2000 µmol m^*-*2^ s^*-*1^, all of these studies applied the high light treatment to low light-adapted plants and focused on the response, or acclimation, to the high light [18, 20, 70, 71, 72, 73, 74, 75, 76]. We, on the other hand, focused on the steady-state transcriptional activity in the four species we examined grown from the seedling stage to maturity under either low or high light.

We have shown that between 68.9 and 85.3% of genes from the four species were not differentially expressed between the light treatments (Table 1). Furthermore, after performing homology grouping and integrating its results with gene differential expression analysis we identified a total of 44,663 HGs containing genes that did not respond to the treatment (Table 2), as well as 631 out of the total 10,352 CHGs containing genes that have the same response to the treatment in all species (Figure 3). None of these genes can, therefore, cause the higher photosynthetic LUE under high light of *H. incana*, and were therefore not considered in our further analysis. We thus believe that the combination of magnitude and application of treatment in our study gives us a sizeable “power of elimination” when dealing with complex transcriptomic datasets.

### Dealing with the complexity of across-species transcriptomic comparisons

The limited set of studies comparing the transcriptomes of different plant species [77, 78, 79, 80, 81] is proof of the novelty of between-species comparative transcriptomics. In contrast to previous studies, we used a panproteome built with PanTools [82] to infer gene homology relationships. We made use of optimised homology grouping, based on the organization of universal single-copy orthologs (BUSCO gene sets, [83, 84]), that is unique to PanTools [44, 45]. This method determines the optimal strictness of protein-clustering settings, given the phylogenetic distance between the proteomes in the data set.

Integrating HGs with per-species transcript abundance and differential expression data presented us with the challenge of comparing transcript abundances across species. Canonical normalization methods, such as the transcripts-per-million (TPM) normalization we used in our study, do not yield abundance measures that can be compared between species [53]. In the absence of a widely accepted approach to compare normalised transcript abundances across species, we decided to estimate the transcriptional “baseline” of the four species. We extracted expression data for all the non-differentially expressed (non-DE) genes belonging to single-copy core HGs and calculating gene-by-gene log_2_-ratios between transcript abundances. Inspection of the distributions of these ratios revealed that *A. thaliana* has on average a slightly higher transcriptional baseline than *B. rapa* and *H. incana*, which in turn have a slightly higher baseline than *B. nigra*. We decided to control for these differences when comparing transcript abundance across species by calculating pairwise log_2_-ratios between TPM-normalized transcript counts and relating them to the ratios calculated for non-DE genes. As we have shown for the genes highlighted in the pathway analyses, the differences between TPM counts are much larger than what could be explained by differences in “baseline” transcription (Figures S7, S9, S10, S11, S12), and therefore have biological meaning.

### Across-species comparison of differential gene expression highlights differences in photosynthetic pathways

The analysis of differential gene expression we performed individually on all four species in this study revealed similar percentages of differentially expressed (DE) genes for *thaliana, B. rapa*, and *H. incana*. Indeed, the cumulative percentage of DE genes in response to HL ranged between 23% and 31% (Table 1). This is in line with previous studies reporting that roughly 20% of the *A. thaliana* transcriptome is responsive to light [85, 71]. For *nigra*, on the other hand, only about 15% of the genes were DE under high light. We do not believe that this difference has a biological explanation, but that it is the result of the very large number of gene models included in the *B. nigra* annotation. Many of these gene models are likely to be annotation artifacts rather than actual genes, as shown by the large number of *B. nigra* genes clustering separately from genes of the other species in the panproteome (Figures 2, S3).

After quantifying gene expression for our four species and inferring homology relationships between genes, we performed a number of untargeted analyses aimed at giving us a non-biased overview of the biological processes and pathways most affected when comparing the transcriptomes from the LL and HL treatments. By making use of the panproteome, we were able to perform these analyses on specific groups of genes, namely the core homology groups containing DE genes in all four species combined, the Brassiceae species (*B. rapa, B. nigra*) and *H. incana*) as a group, and *H. incana* on its own. Considering the higher photosynthesis rates we previously reported for the Brassiceae species [21], one might expect results linked to photosnynthetic LUE to come from the HGs showing differential expression for the Brassiceae species, or from the HGs with genes showing DE in *H. incana* alone. Nevertheless, the most promising results came from enrichment analyses on the HGs containing genes deferentially expressed across all four species. Indeed, out of a total of nine KEGG pathways enriched in this kind of HGs, three pathways mentioned photosynthesis in their name.

One striking finding of our targeted analysis of the three photosynthesis-related pathways was that most of the associated genes appeared to either have lower or unchanged expression under the HL treatment. This was expected for the “Photosynthesis - antenna proteins” pathway (ath00196), including all photosystem antenna genes, based on experimental evidence that plants growing under high light will reduce the size of their antennas [86]. However, this trend of lower or unchanged gene expression was unexpected for genes related to photosynthetic light reactions (included in the “Photosynthesis” pathway, ath00195) and carbon metabolism (included in the homonymous pathway, ath01200). Recent studies of changes in the *A. thaliana* proteome in response to irradiance increase or switch from controlled to field conditions have highlighted increases in abundance for most proteins involved in light reactions [87, 60]. Furthermore, ample experimental evidence has been collected in the past showing that plants acclimating to high light develop a higher carbon fixation metabolism via increased protein levels [88]. A few considerations arise from the discrepancy between this evidence and the results of our transcriptome analysis. The first is that, as already discussed above, previous studies focused on acclimation responses to higher light, while ours was conducted on plants that grew under constant high or low light, and therefore the transcriptome snapshot obtained in our study might represents a much different gene and protein regulation situation than what was previously studied. Furthermore, it is important to point out that while transcriptome analysis highlights genes that are potentially involved in high photosynthetic LUE, it cannot inform us on downstream proteome dynamics. Thus, we currently cannot say whether higher gene expression is a consequence of higher protein turnover due to e.g. photodamage, or if it enables for higher protein abundance, thus potentially enabling for higher biochemical capacity in the photosynthetic reactions. The opposite is naturally true for lower gene expression, and therefore this analysis does not allow us to conclude whether that is the result of higher protein stability or lower protein abundance requirements.

Finally, our in-depth analysis of gene expression for three KEGG pathways revealed that differential gene expression is only one of the ways *H. incana* achieves higher transcript abundances, potentially enabling its higher photosynthetic light-use efficiency. While we identified four genes encoding photosystem subunits (*PSBD, PSBE, PSAA, PSAJ*) whose transcript levels were significantly higher in *H. incana* plants grown under HL, we identified other genes such as *LHCB8* and *LHCA6* having a striking transcript abundance in *H. incana* plants grown under both irradiances. This appears to be achieved in two additional ways: while the *LHCB8* gene is present in a single copy in *H. incana* and all other species, and the abundance of its transcript in *H. incana* can be explained with a constitutive overexpression of the gene, the *LHCA6* gene is present in three copies in *H. incana* as opposed as the single copy of the other three species. Each *LHCA6* copy is expressed in *H. incana* at levels that appear to be slightly higher than those of other species, but the cumulative expression of the three copies results in a substantially higher transcript abundance for the gene. These strategies to achieve higher gene expression form an interesting lead to further investigate the precise relationship between expression levels, protein abundance and turnover, and ultimately the photosynthetic light-use efficiency of *H. incana*.

### Possibilities to explore other processes related to photosynthesis

While we decided to limit our research to KEGG photosynthesis pathways, we acknowledge that photosynthesis is a highly complex process involving other key pathways. We hypothesize that genes involved in transpiration, heat dissipation, stress response, and nutrient uptake and cycling will play a role in supporting higher photosynthetic efficiency. While previous studies identified transcriptional responses to high irradiance connected to heat-shock response [18, 76], ribosome biogenesis and transcriptional activity [76], lipid remodeling [18], flavonoid biosynthesis [74, 20], a comprehensive picture of these responses is still far from being available [89]. Based on what emerged from our targeted analysis on photosynthetic pathways, approaching our dataset in a different way than via enrichment analysis will likely reveal how the these processes are playing a role in high-light photosynthesis. Our resource will therefore provide means to further explore the genetic basis of high photosynthetic efficiency under high light.

### Prospects for future research

In this study, we have highlighted three different strategies that *H. incana* can employ to achieve higher transcript abundances for genes that potentially play a key role in its photosynthetic efficiency. Given that our analysis pipeline allows the retrieval of TPM-normalized counts for all expressed genes in each of the employed species, and that we established a method to estimate baseline differences in transcript abundances for the various species, an additional study of transcript abundances irrespective of differential expression might provide further clues on the mechanisms allowing *H. incana* to achieve higher photosynthetic light-use efficiency. Despite showing that most genes involved in photosynthetic reactions have lower expression as a response to high light in all analysed species, we have identified a number of genes that are either highly expressed in response to high light or have a constitutive higher expression in *H. incana*. Of these genes, *LHCB8* and *LHCA6* appear as very promising targets for further analysis, as the function of the first is still unclear and the higher expression in *H. incana* of the second cannot be explained with current literature.

It is important to stress once more how this experiment aimed at obtaining a snapshot of the operation of high photosynthesis rates, rather than at their establishment during leaf development. While our experiment uncovered some genes that might be playing a role in supporting high photosynthetic activity under high irradiance, future transcriptomics investigations on time series collected throughout leaf development will be crucial to understand which genes and processes enable the establishment of high photosynthetic light-use efficiency.

## Conclusions

This study provides an analysis of the transcriptomes of *A. thaliana, B. rapa, B. nigra*, and *H. incana* plants grown under constant low and high irradiance, rather than the acclimation response to high irradiance. By combining gene expression quantification and differential expression analysis with a panproteome-based homology grouping, we quickly and efficiently identified expression patterns shared by the various species, or unique to one of them. Following an untargeted approach, we observed an enrichment for genes involved in photosynthetic pathways. A closer look at the expression of all genes belonging to these pathways allowed us to reveal that in comparison to other Brassicaceae species, *H. incana* growing under a high light treatment achieves higher expression of genes related to photosynthesis via three different modes: “canonical” differential expression between low and high light, constitutive higher expression of single-copy genes, or cumulative higher expression obtained by simultaneous expression of multiple gene copies. Besides identifying genes such as *LHCB8* and *LHCA6*, whose higher expression in *H. incana* growing under high light prompts for a detailed investigation of their role in photosynthetic LUE under high irradiance, we believe that analysing the genes undergoing differential expression specifically in *H. incana* will further clarify the role of non-strictly photosynthetic genes in supporting the species’ striking photosynthetic performance. Therefore, we expect the resource we established with this study to provide further, extensive knowledge on the genetic strategy employed by *H. incana* to support its high photosynthetic light-use efficiency.

## Data Availability

Raw sequencing data has been deposited to NCBI and can be found under BioProject PRJNA1001172. All scripts used for data analysis are available on the 4TU.ResearchData platform with the DOI 10.4121/5b88cdf2-eb5f-4033-8ece-1f3f488a1f83.v1

## Acknowledgements

We thank Ludovico Caracciolo for his help during the building of the growth setup and for helping us with plant growth and sampling. We thank Nam Hoang for the critical discussions on the methodology and results of this study, as well as for helping us with plant sampling. Furthermore, we thank Vittoria Clapero for her involvement in the early stages of this study, Robin van Esch and Dirk-Jan van Workum for their support for parts of the analysis, and Eric Schranz for his useful contributions in discussions on this study.

## Funding

F.G. and S.B.L are supported by a private grant.

## Conflict of interest statement

None declared.

Garassino *et al*. – page 13

## Author contributions

F.G. and S.S. designed the study in consultation with M.A. and J.H.. F.G. grew and sampled plants. F.R.M. extracted and evaluated RNA from all samples. T.C. performed preliminary transcriptome analyses, which F.G. extended and finalized with help from S.B.L.. H.N. and S.S supervised transcriptome analysis. F.G., S.B.L., H.N., and S.S. were involved in writing the manuscript. M.A. and J.H. provided guidance on the biological interpretation of results and revised the manuscript. All authors have approved the contents of the manuscript.

## Notes

https://www.ncbi.nlm.nih.gov/bioproject/?term=PRJNA1001172

https://doi.org/10.4121/5b88cdf2-eb5f-4033-8ece-1f3f488a1f83.v1

### Competing Interest Statement

The authors have declared no competing interest.

