## Supplementary Figures for "Comparative transcriptomics of Hirschfeldia incana and relatives highlights differences in photosynthetic pathways"

### Supplementary Figures, belonging to Garassino et al. (2023), "Comparative transcriptomics of Hirschfeldia incana and relatives highlights differences in photosynthetic pathways"

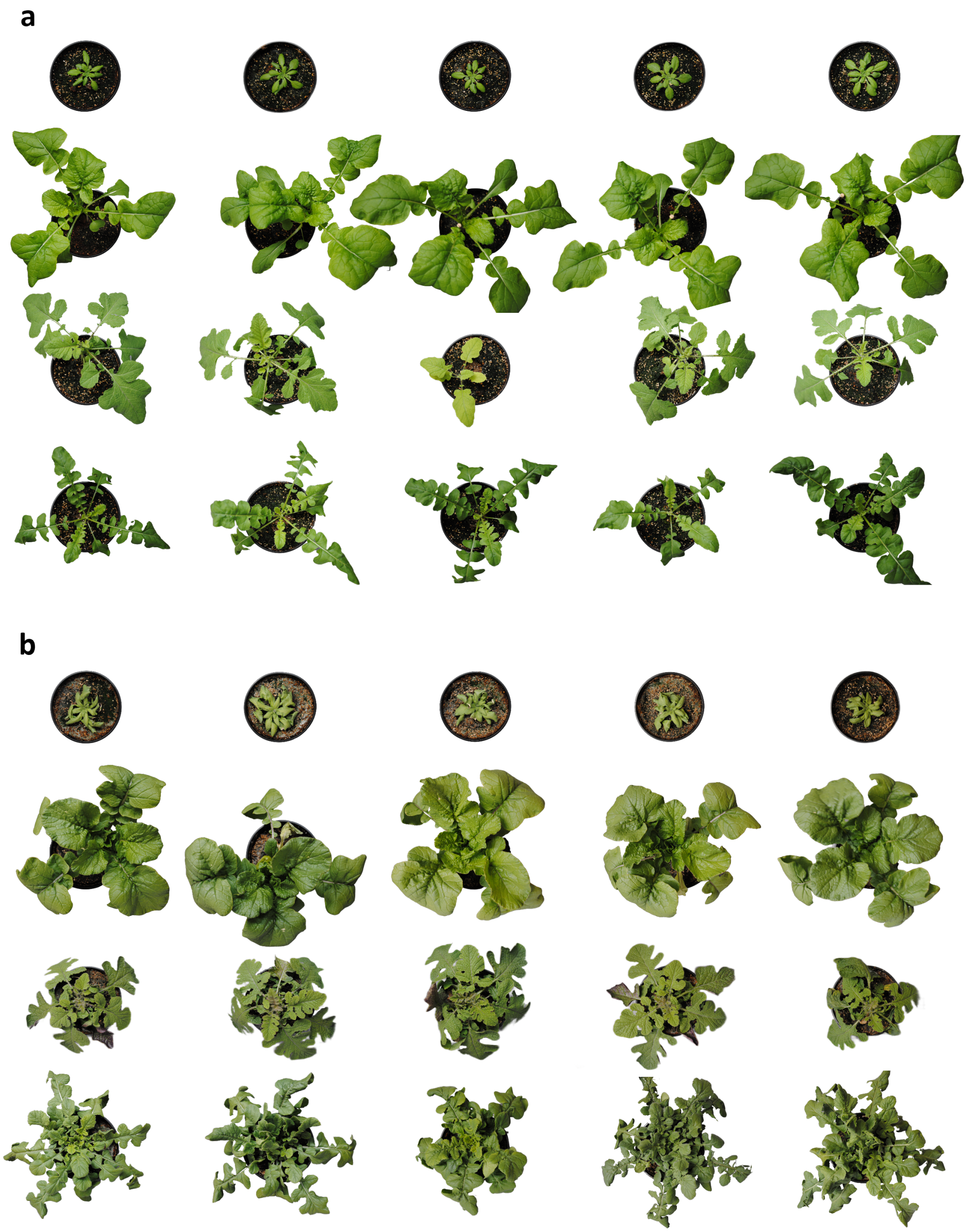


Figure S1: Pictures of all plants employed in this study, grown under low (a) or high light (b). For both sets, from top to bottom: *A. thaliana*, *B. rapa*, *B. nigra*, and *H. incana* plants.

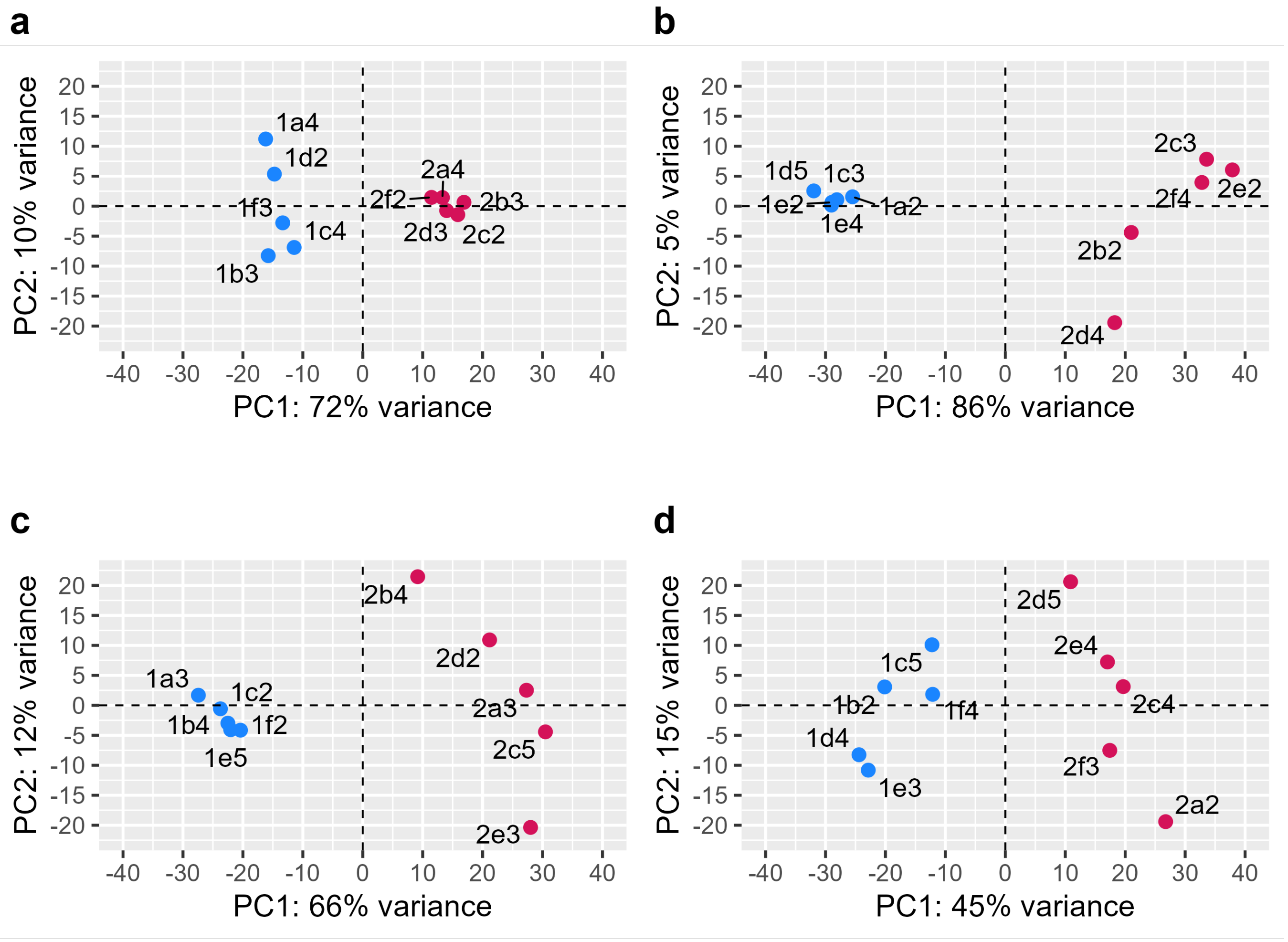


Figure S2: Per-species principal Component Analysis (PCA) plots of transcript counts transformed by means of regularized logarithms. (a) *A. thaliana*, (b) *B. rapa*, (c) *B. nigra*, (d) *H. incana*. Blue dots represent samples from plants grown under low light, and red dots samples from plants grown under high light.

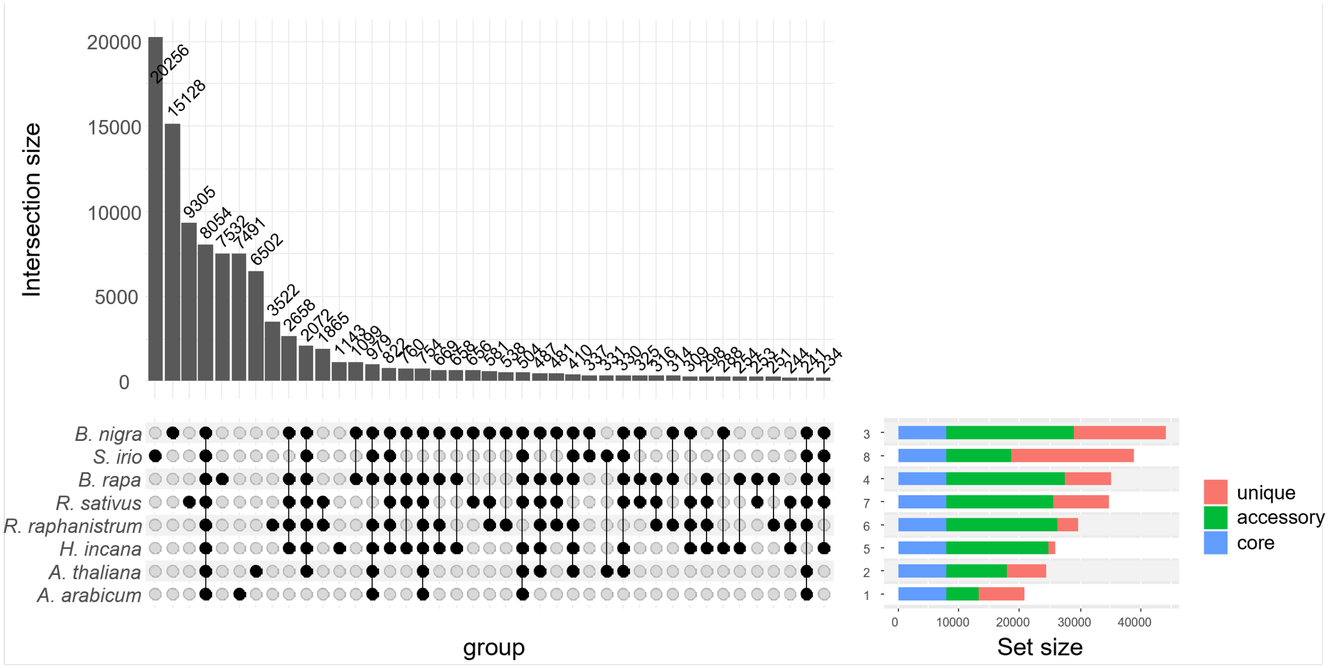


Figure S3: UpSet plot for the whole panproteome. Vertical bars represent the number of homology groups, classified by presence/absence of genes from the various species as illustrated at the bottom of the figure. Horizontal colored bars show how many of the groups containing at least one gene from a species fall within one of three categories: core groups, i.e. those containing at least one gene from all four species; accessory groups, i.e. those containing genes from more than one species but less than all species; and unique groups, i.e. those containing only genes from a single species.

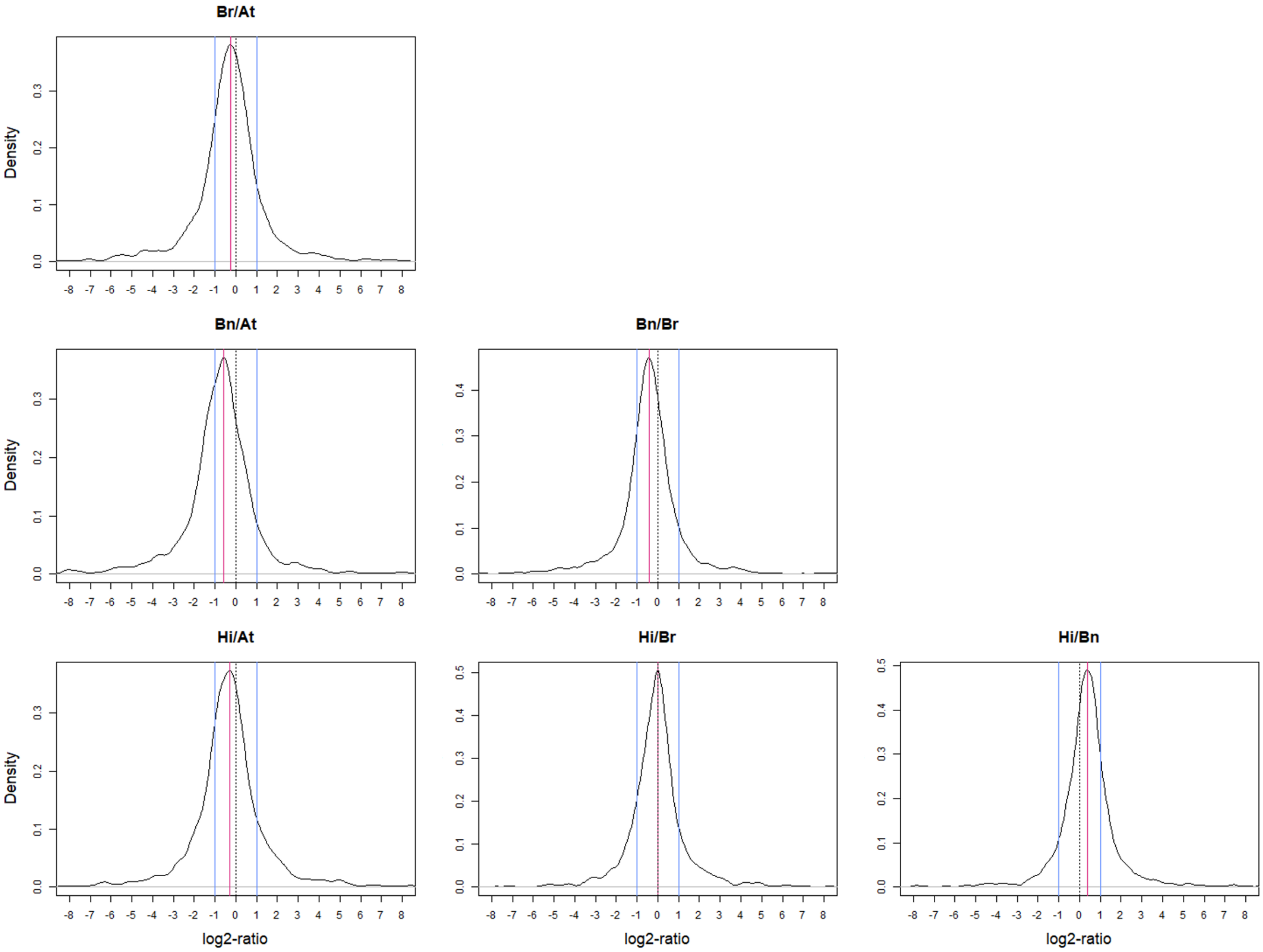


Figure S4: Distribution of log2-ratios between TPM-normalised transcript abundances for 1,692 single-copy non-DE orthologs. The black dotted line (log2-ratio of 0) represents a TPM-ratio of 1, the blue lines represent TPM-ratios of -2 (left) and 2 (right), and the red line corresponds to the peak of the distribution. At: *Arabidopsis thaliana*, Br: *Brassica rapa*, Bn: *Brassica nigra*, Hi: *Hirschfeldia incana*. Values for areas under the curve (AUCs) and peak coordinates can be found in Table S9. The ratios shown in this figure are specular to those in the plots not shown (e.g. At/Br).

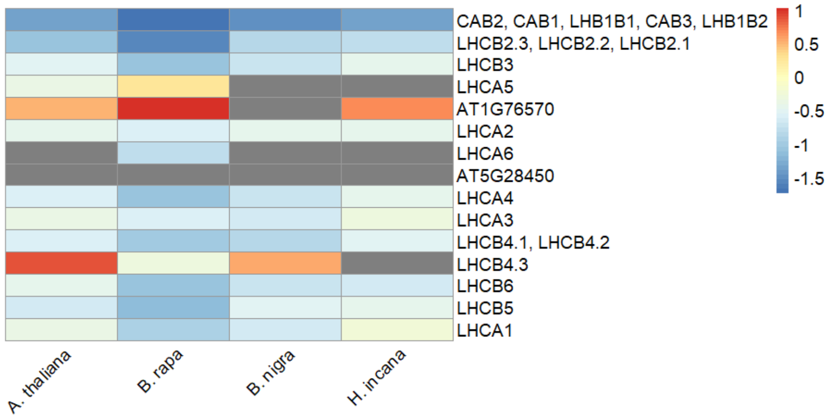


Figure S5: Heatmap of log2-fold changes of nuclear and chloroplast genes associated to KEGG pathway "Photosynthesis - antenna proteins" (ath00196). Averages were calculated if multiple genes were present in an orthology group for one species. Grey fields indicate that no significant log2-fold change was present for the genes of the corresponding species belonging to a specific orthology group.

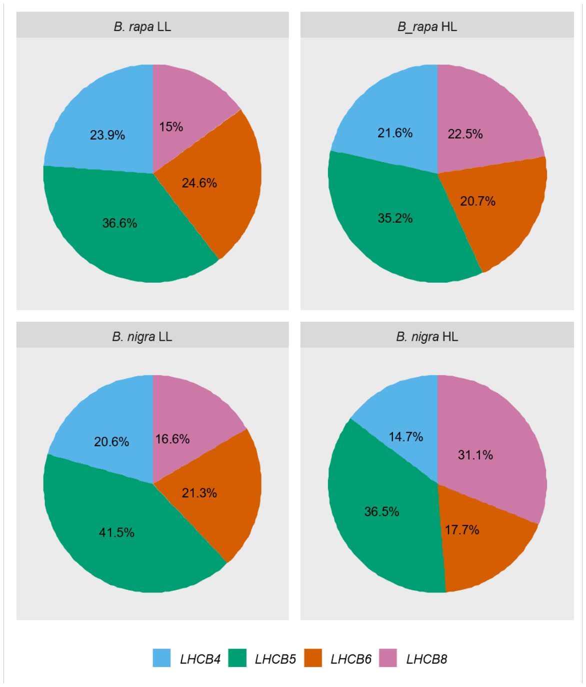


Figure S6: Pie charts representing the relative abundance of transcripts originating from genes encoding components of PSII minor antenna for *B. rapa* and *B. nigra*.

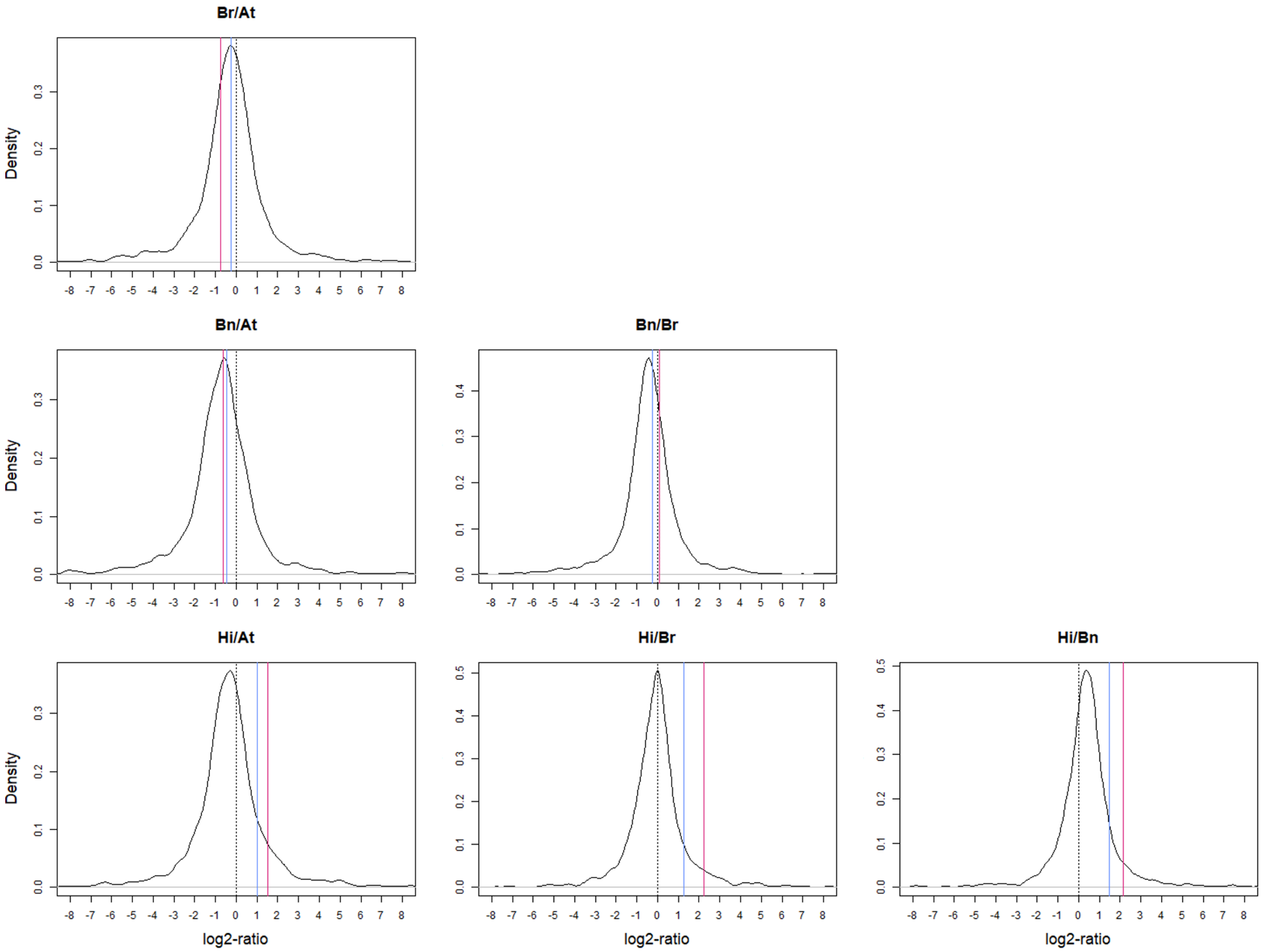


Figure S7: Positioning of pairwise log2-ratios between TPM-normalised transcript abundances for *LHCA6* transcripts, relative to the distribution of pairwise ratios for the 1,692 single-copy "non-responsive" orthologs. The blue line represents the ratio for counts in plants grown under the LL treatment, while the red line the one for counts in plants grown under the HL treatment. The AUCs to the left and right of the *LHCA6* ratios can be found in Table S15.

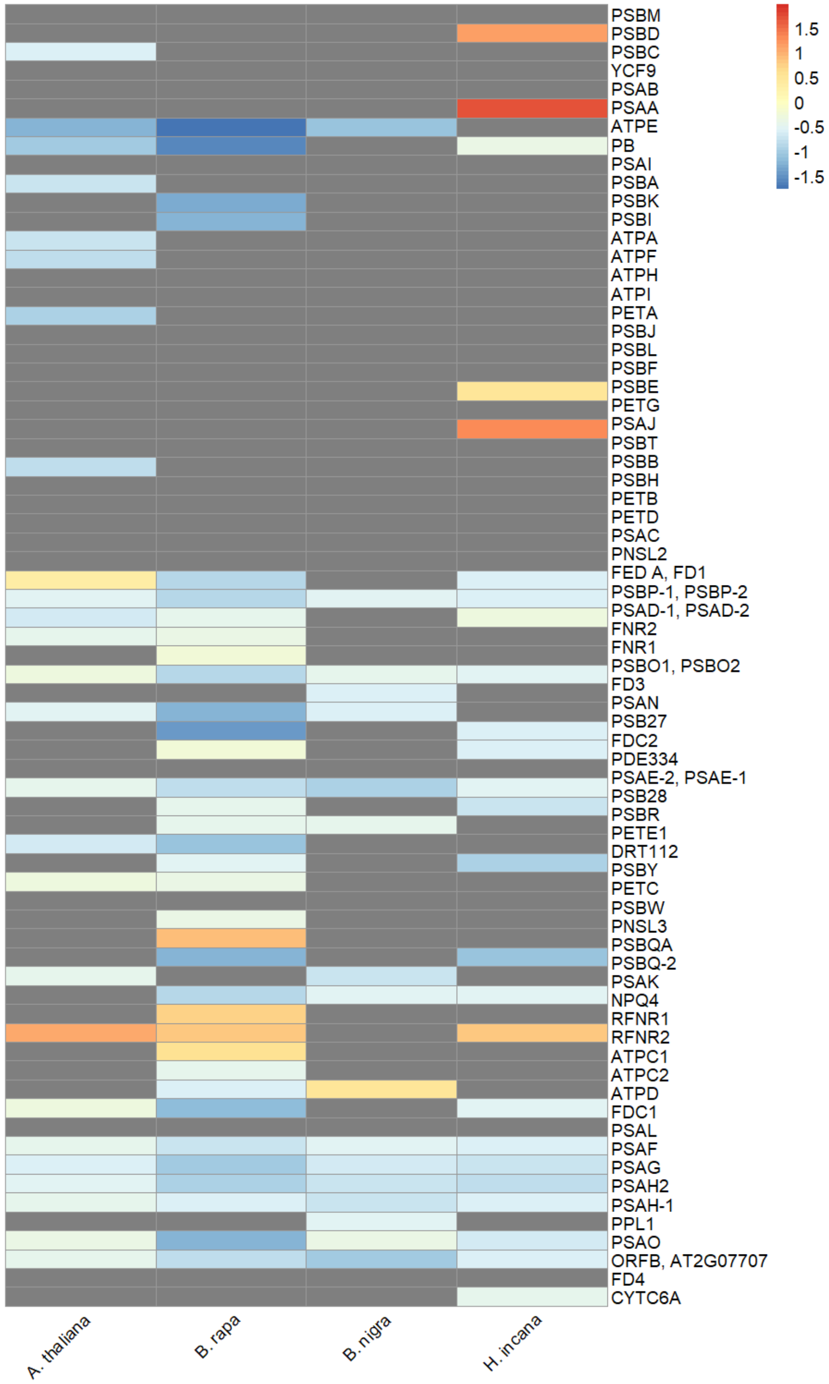


Figure S8: Heatmap of log2-fold changes of nuclear and chloroplast genes associated to KEGG pathway "Photosynthesis" (ath00195). Averages were calculated if multiple genes were present in an orthology group for one species. Grey fields indicate that no significant log2-fold change was present for the genes of the corresponding species belonging to a specific orthology group.

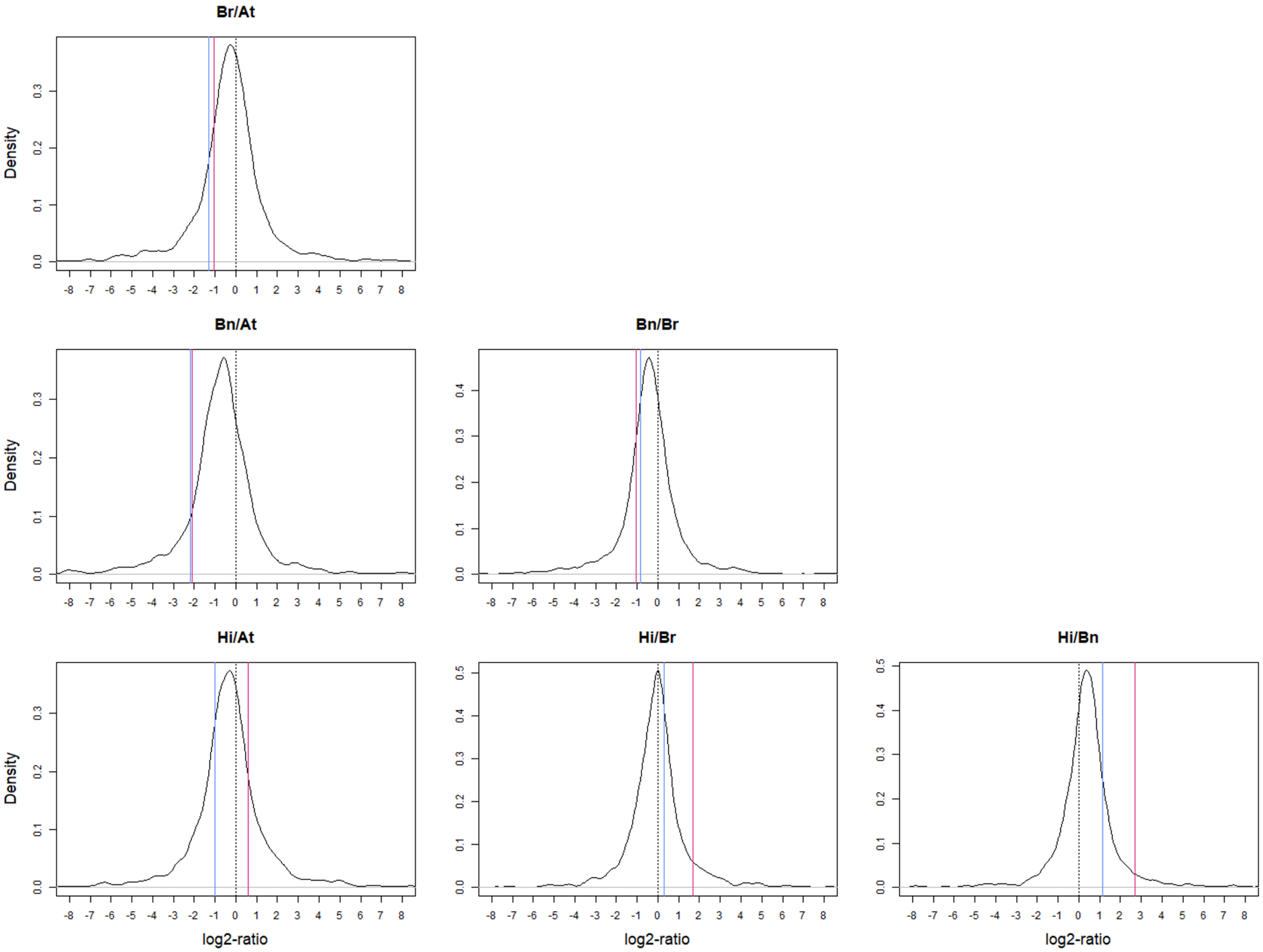


Figure S9: Positioning of pairwise log2-ratios between TPM-normalised transcript abundances for *PSBD* transcripts, relative to the distribution of pairwise ratios for the 1,692 single-copy "non-responsive" orthologs. The blue line represents the ratio for counts in plants grown under the LL treatment, while the red line the one for counts in plants grown under the HL treatment. The AUCs to the left and right of the *PSBD* ratios can be found in Table S16.

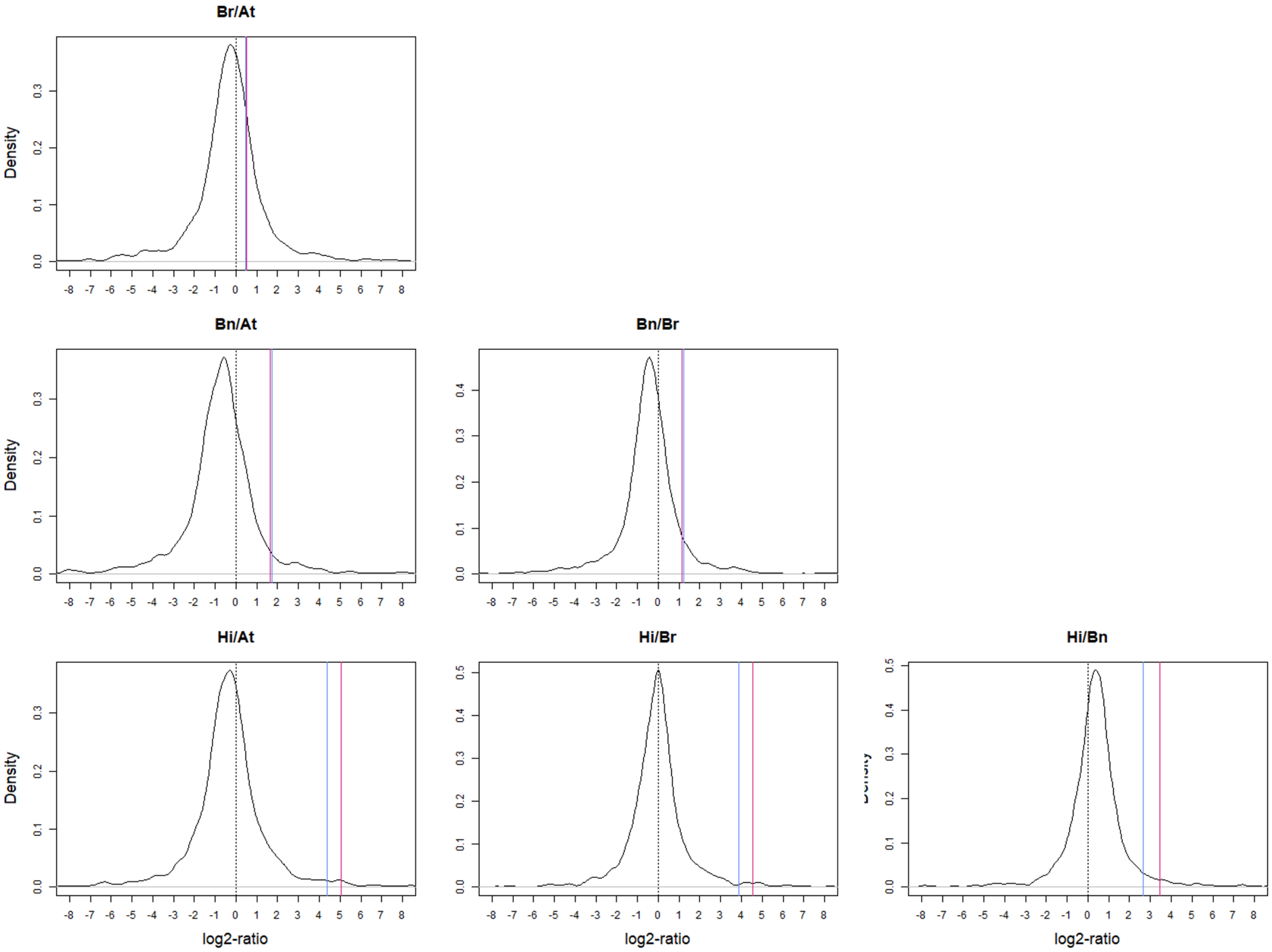


Figure S10: Positioning of pairwise log2-ratios between TPM-normalised transcript abundances for *PSBE* transcripts, relative to the distribution of pairwise ratios for the 1,692 single-copy "non-responsive" orthologs. The blue line represents the ratio for counts in plants grown under the LL treatment, while the red line the one for counts in plants grown under the HL treatment. The AUCs to the left and right of the *PSBE* ratios can be found in Table S17.

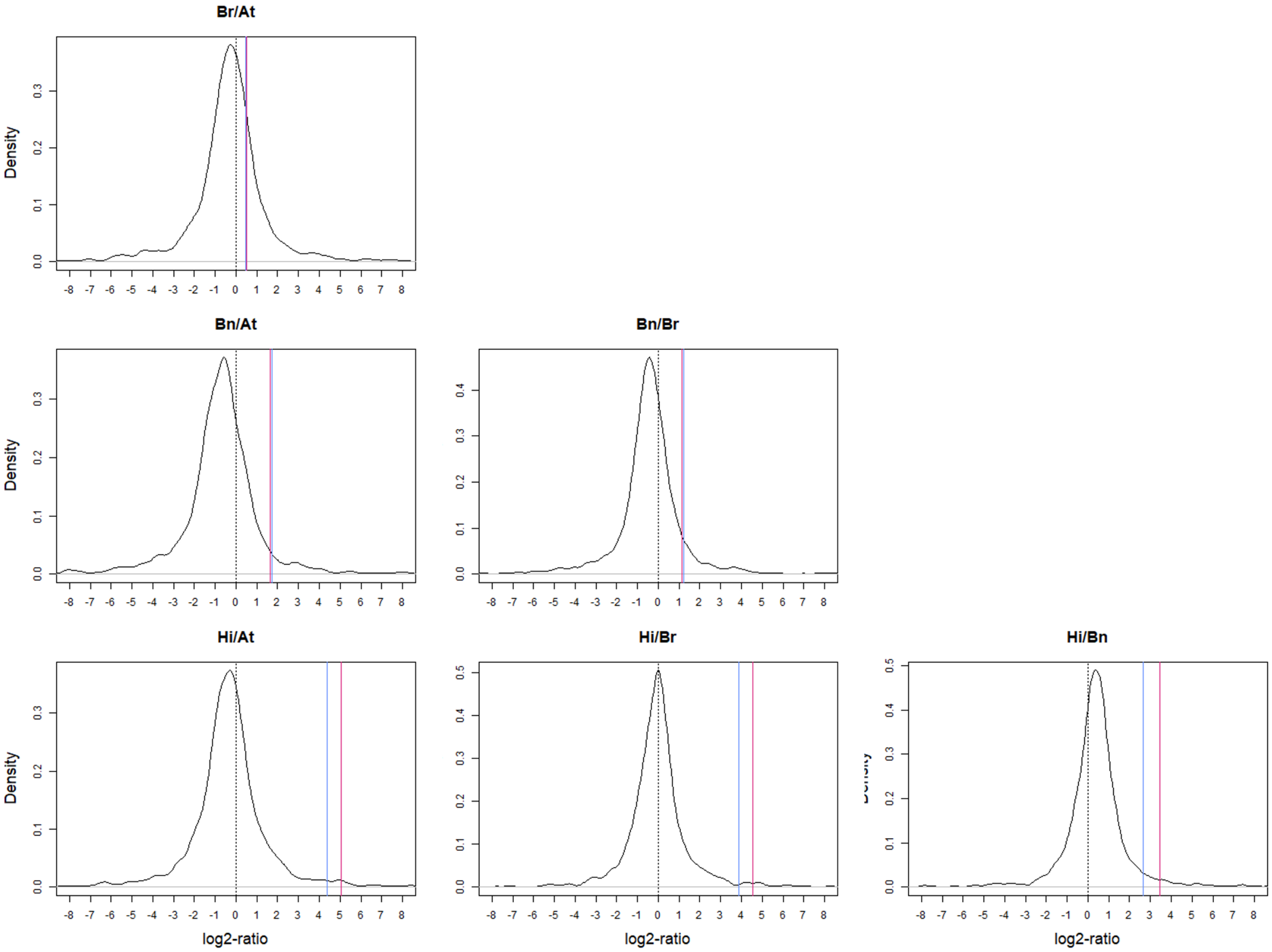


Figure S11: Positioning of pairwise log2-ratios between TPM-normalised transcript abundances for *PSAA* transcripts, relative to the distribution of pairwise ratios for the 1,692 single-copy "non-responsive" orthologs. The blue line represents the ratio for counts in plants grown under the LL treatment, while the red line the one for counts in plants grown under the HL treatment. The AUCs to the left and right of the *PSAA* ratios can be found in Table S18.

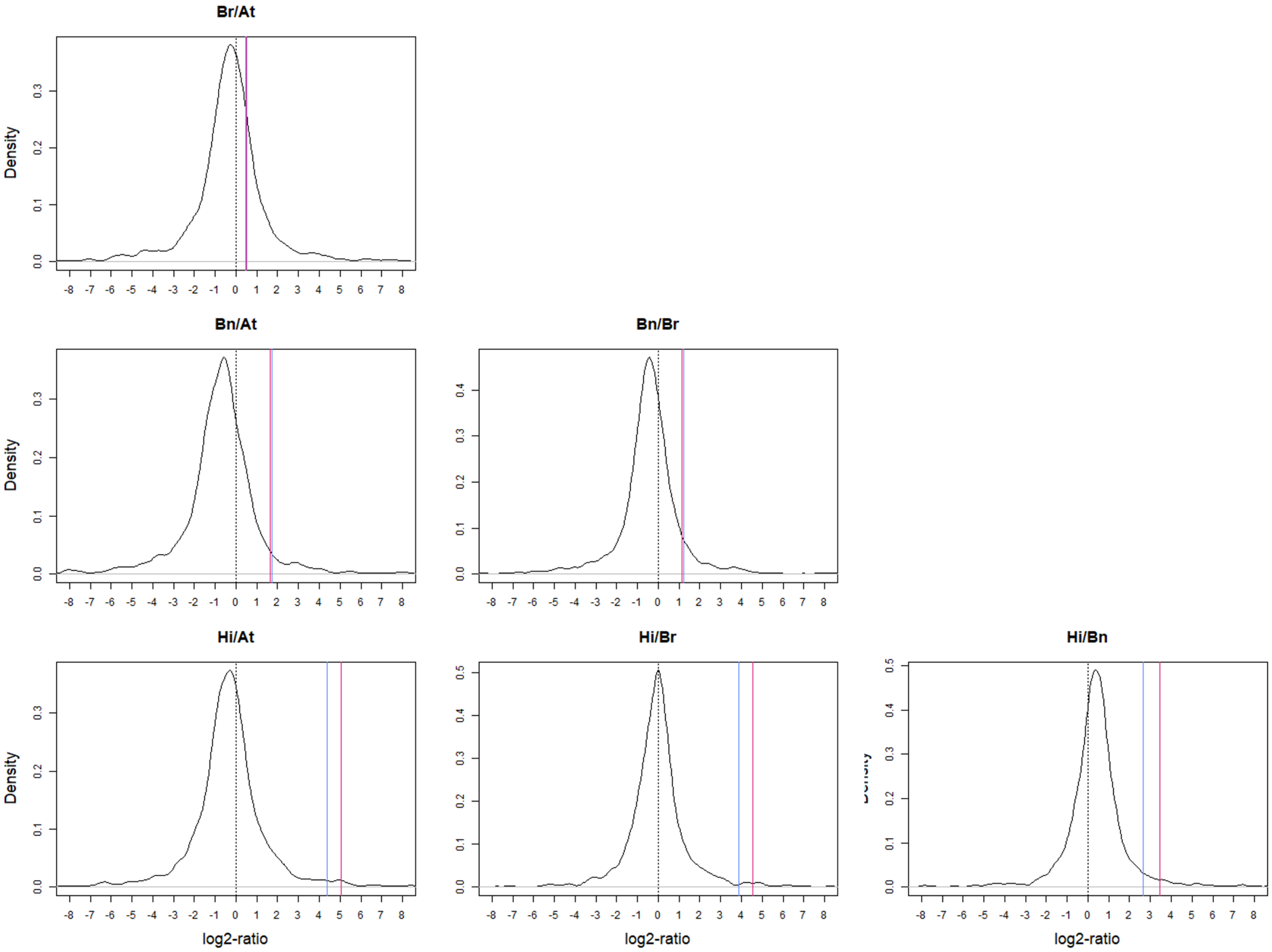


Figure S12: Positioning of pairwise log2-ratios between TPM-normalised transcript abundances for *PSAJ* transcripts, relative to the distribution of pairwise ratios for the 1,692 single-copy "non-responsive" orthologs. The blue line represents the ratio for counts in plants grown under the LL treatment, while the red line the one for counts in plants grown under the HL treatment. The AUCs to the left and right of the *PSAJ* ratios can be found in Table S19.

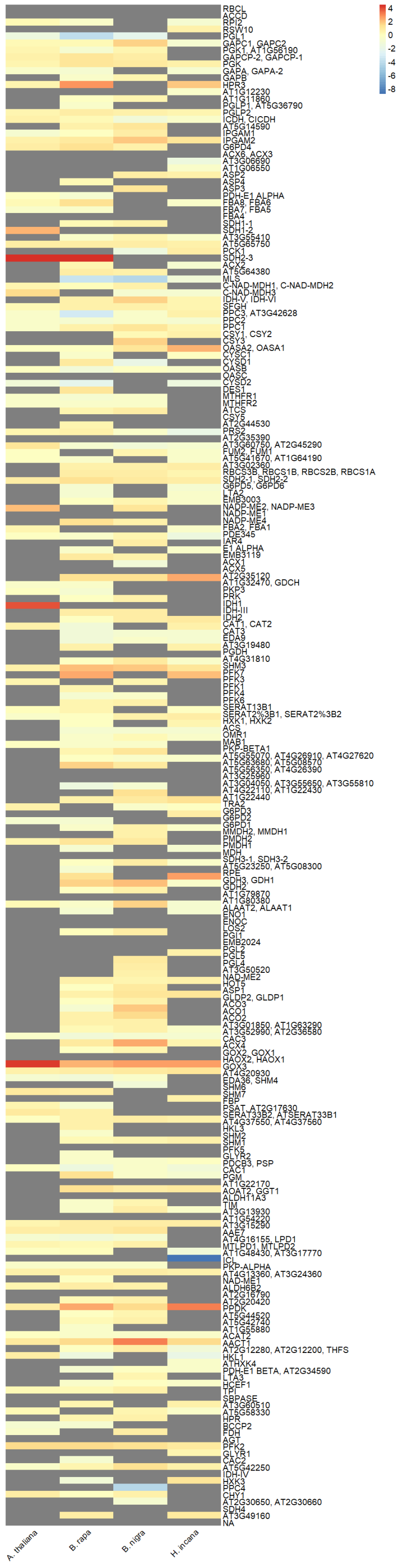


Figure S13: Heatmap of log2-fold changes of nuclear and chloroplast genes associated to KEGG pathway "Carbon metabolism" (ath01200). Averages were calculated if multiple genes were present in an orthology group for one species. Grey fields indicate that no significant log2-fold change was present for the genes of the corresponding species belonging to a specific orthology group.

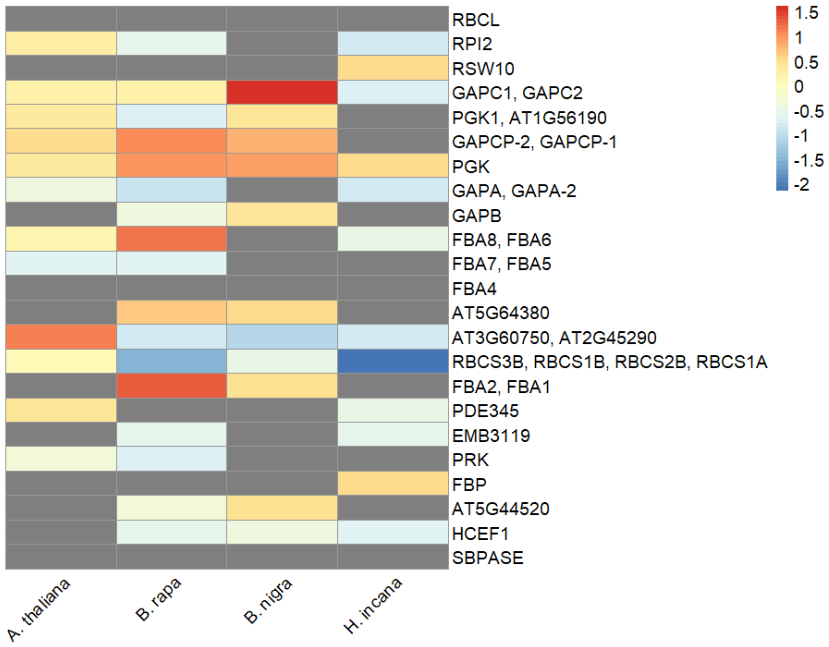


Figure S14: Heatmap of log2-fold changes of nuclear and chloroplast genes associated to KEGG module "Reductive pentose phosphate cycle (Calvin cycle)" (M00165). Averages were calculated if multiple genes were present in an orthology group for one species. Grey fields indicate that no significant log2-fold change was present for the genes of the corresponding species belonging to a specific orthology group.
